# Population activity of mossy fibre axon input to the cerebellar cortex during behaviours

**DOI:** 10.1101/2025.03.23.644738

**Authors:** Hana Roš, Yizhou Xie, Sadra Sadeh, R. Angus Silver

## Abstract

The cerebellum gathers information from the neocortex via the cortico-ponto-cerebellar pathway, and forms sensorimotor associations for predicting movements and their sensory consequences. However, little is known about the functional properties of this major input to the cerebellar cortex. Recordings from individual cerebellar mossy fibre axons (MFAs) have shown that they convey sensory and motor information, but nothing is known about their population code. Here, we report that the population activity of pontine MFAs is heterogeneous, high-dimensional and that different subpopulations of MFAs are active during quiet and active behavioural states. Population activity occupied a substantial fraction of the state space and some MFAs are particularly informative about behaviour. Surprisingly, positively and negatively modulated MFAs are intermingled, suggesting granule cells integrate opposite-signed inputs to generate mixed bidirectional sensorimotor representations. Our results establish that neocortex and cerebellum can communicate with a low redundancy, high capacity, bidirectional population code, which is well-suited for forming sensorimotor associations.

**Highlights:**

● Population activity of main mossy fibre axonal input to cerebellar cortex.
● Ponto-cerebellar code is high dimensional during behaviours.
● Behavioural information conveyed by bidirectional population code.
● Modelling predicts heterogeneous response properties of granule cells.

## Introduction

The brain gathers information about the body and the surrounding world, enabling it to build internal representations and to plan and execute movement. A key role of the cerebellum is to form associations that predict actions and their sensory consequences^1,2^. Generating such internal models enables smooth, accurate movements by circumventing slow sensory and proprioceptive feedback^3^. Learning to predict the sensory consequences of movements is important for active sensing and distinguishing between self-generated and external sensory input^4,2^. The sensorimotor cortico-ponto-cerebellar loop^5,6^ is therefore critical for interacting with the environment and adapting to changing conditions. However, the nature of the sensorimotor computations carried out in the cerebellum remain uncertain, due in large part to a lack of information on the properties of the synaptic input to the cerebellar cortex.

Mossy fibre axons (MFAs) originate from multiple sources and are known to convey sensory and motor information to the input layer of the cerebellar cortex. The largest source of mossy fibres is from the basal pontine nucleus (BPN)^7,8^, which forms part of the cortico-ponto-cerebellar pathway^9^. The BPN is predominantly innervated by projections from sensory, motor and prefrontal regions of the neocortex^6,10,11,12,13^ but also receives excitatory feedback from the cerebellar nuclei^14^ suggesting that the BPN performs active processing, rather than acting as a simple relay of neocortical output^7,8,15,16^.

MFAs formed by neurons in BPN project widely across the cerebellar cortex, with the hemispheric regions, such as CrusI/II, receiving the most dense innervation^10,7,9^. Individual BPN MFAs innervate multiple cerebellar lobules ^17^ forming many large presynaptic mossy fibre boutons (MFBs, also referred to as synaptic rosettes^18^. Each synaptic rosette is thought to innervate 12-50 granule cells^19,20–22^ with each granule cell (GrC) receiving input from 4 different MFAs on average^23,106^. This sparse synaptic connectivity onto GrCs and vast neuronal expansion is thought to mix different streams of information^24,25,26,27,28^, sparsen^19^, decorrelate^29^, and support high dimensional representations^29,24^, thereby separating overlapping MFA activity patterns^30^, prior to downstream associative learning^31,29,32,33,34^. Recent experimental recordings from GrCs have revealed a considerably higher level of activity ^35,36,37^ than the ultra-sparse population codes predicted by classical models^33,34^. Moreover, such GrC population activity can support dense low dimensional representations of simple and learned behaviours^35,36,38^ or distributed high dimensional population representations of more complex spontaneous behaviours^24^. However, it is not known how these representations are generated, because in contrast to GrCs, little is known about the properties of the cortico-pontine MFA input to the cerebellar cortex during behaviour.

Early single unit recordings from putative MFAs in monkey cerebellum suggested that they convey rate-coded information on limb and digit kinematics^39^. These findings have been confirmed using patch-clamp recordings from a small number of individual MFAs in awake mice, which show that their firing encodes information on limb trajectory^40^. Recordings from MFAs in anaesthetised mice have shown that they respond to air puffs by firing at very high frequencies ^41^ and that MFAs in the vestibular cerebellum linearly encode angular head velocity^42^. Sustained signalling of MFAs has also been observed with postsynaptic recordings of EPSCs from GrCs in awake mice, which have revealed that MFA synaptic input is significant during quiescent periods and further elevated during active whisking, suggesting that MFAs encode both whisker position/set point and kinematics^43^. The finding that individual MFAs sustain signalling *in vivo* over long periods fits with results from acute slices, which show that MFA-GrC synapses can sustain firing over a wide frequency bandwidth through both presynaptic^44,45,46^ and postsynaptic^27,47,7,8,48^ specialisations. While these studies confirm that individual MFAs are well adapted to continuously update the cerebellar cortex with sensory and motor information, little is known about how they represent such information at the population level within the input layer of the cerebellar cortex. The lack of data on MFA population activity therefore represents a major gap in our understanding of cerebellar function.

To address this knowledge gap, we imaged the activity of identified populations of MFAs in the GrC layer of mouse cerebellar cortex using Acousto-Optic Lens (AOL) 3D two-photon microscopy^49^. Our results reveal that the population activity of MFAs arising from the BPN is heterogeneous, with different subpopulations of MFAs increasing and decreasing their activity during locomotion and whisking. The MFA population activity was high dimensional and formed orthogonal subspaces during quiet and active spontaneous behaviours. Surprisingly, MFAs of opposite sign were intermingled within local regions, raising the possibility that a single GrC receives mixed combinations of these inputs. The high dimensional, bidirectional MFA population code appears well suited for continuously updating the cerebellar cortex with sensory, motor and cognitive states, information that is crucial for forming internal models that enable coordinated action and predictions of their sensory consequences.

## Results

### Heterogeneous pontine mossy fibre population activity during spontaneous behaviours

To investigate population-level activity in identified MFAs (Figure 1A) we injected AAV9-CAG-Flex-GCaMP6f in Slc17a7-IRES-Cre transgenic mice to restrict the extent of expression to the BPN^17^ (S1A). Specific expression of GCaMP6f in BPN MFAs and their distribution across Crus I/II of the cerebellar cortex was confirmed *post hoc* with confocal imaging of fixed tissue (Figure 1B and S1B,C). To monitor the activity of as many MF synaptic varicosities (referred to as MFBs) as possible in Crus I/II, we formed a cranial window over these areas (Figure 1C and S1D) and used an AOL 3D two-photon microscope to perform multi-plane or multi-patch measurements^49^ in head-fixed mice freely locomoting on a cylindrical treadmill. GCaMP6f expression was dense and widespread, with hundreds of labelled MFBs within each 250 x 250 x 100 µm imaging volume (Figure 1D), consistent with anatomical studies^17^. This strategy enabled us to simultaneously record the activity from an average of 280 ± 5 BPN MFBs per session (mean ± SEM; *n* = 16 sessions) across 4 mice (*N* = 4).

**Figure 1.**
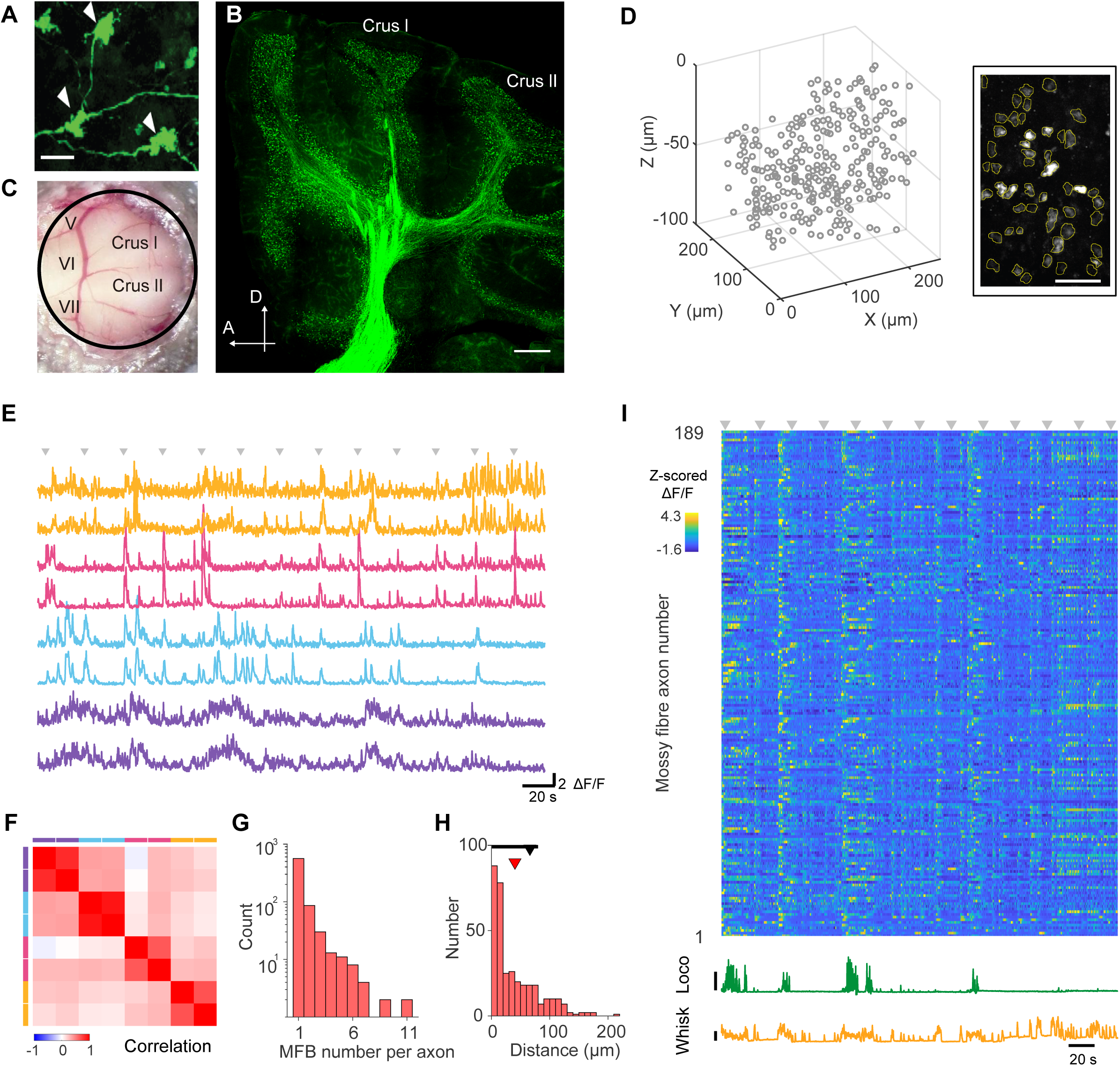
**Population imaging of identified ponto-cerebellar mossy fibre axons** (A) Confocal images of mossy fibre axons arising from basal pontine nuclei (BPN) in fixed tissue; scale bar 10 μm; white arrowheads show mossy fibre boutons (MFBs, varicosities/rosettes). Specific labelling was achieved using Cre-dependent AAVs expressing GCaMP6f that were stereotaxically injected into BPN in Slc17a7-IRES-Cre mice. (B) Termination patterns of GCaMP6f-expressing BPN mossy fibres in the cerebellum; scale bar 100 µm; A, anterior, D, dorsal. (C) Chronic window implant (5 mm) above Crus I/II of mouse cerebellum. (D) Location of MFBs in a 3D two-photon imaging volume. Individual imaged patch (right) with regions of interest around individual MFBs (yellow lines) for extraction of fluorescence traces; scale bar 20 μm. (E) Examples of Z-scored GCaMP6f ΔF/F traces from highly correlated MFBs, with each colour corresponding to those identified as arising from the same axon. Arrowheads show timing of mild airpuffs to the ipsilateral whiskers. (F) Matrix showing correlations between MFB activity in (E). (G) Distribution of the number of MFBs per axon (1076 MFBs in total). (H) Distribution of euclidean distance between MFBs on the same axon (345 pairs, n = 16, N = 4). The red arrow shows the mean intervaricosity distance. The black arrow indicates the mean intervaricosity distance along the axon determined previously in fixed tissue. (I) Z-score of ΔF/F traces (range −1.6 − 4.3, 99th percentile of amplitude distribution) for 189 BPN MF axons during a single recording session.

MFAs branch and form *en passant* synaptic boutons (rosettes) along their length^50^. If boutons belong to the same axon, their activity should be a scaled version of each other, assuming they are in the linear regime of GCaMP6f. Comparison of MFBs across each imaging volume during spontaneous activity and mild air puffs to the whiskers, revealed that some MFBs exhibited very similar activity profiles (Figure 1E), suggesting they arose from the same axon. To identify MFBs from the same axon, we applied a modified version of the procedure developed for grouping boutons on GrC axons^24^. The first criterion for identifying pairs of MFBs arising from the same axon was highly positively correlated activity (Spearman correlation coefficient, 𝐶𝐶_𝑠_, of >99 percentile; Figure 1F). To account for differing levels of amplitude variability in fluorescence traces that could arise from different MFB sizes, variations in GCaMP6f expression, depth of imaging and laser power, we then determined the deviation from linear scaling. For this, we calculated the linear deviation ratio (LDR) by normalising the projection vector (v) by the variance of the distribution of baseline fluorescence projected onto v and rejected pairs with an LDR > 1.5 (Method details, S2). This two-step analysis identified 719 MFAs, of which 561 were single MFBs and 158 had between 2 - 11 MFBs on the same axon (*n* = 16, *N* = 4; Figure 1G). The nearest neighbour Euclidean distance between MFBs on the same axon ranged from 3 µm to 212 µm (Figure 1H) with a mean of 40.3 ± 39.6 µm (mean ± SD, *n* = 16, *N* = 4). This is consistent with (but not directly comparable to) the distance between MFBs measured along the axon in adult mice^50^ (66 ± 55 µm). Averaging the fluorescence time series from MFBs arising from the same axon gave an average of 191 ± 4 MFA recordings per session (*n* = 16 sessions, *N* = 4). Comparison of the normalised fluorescence transients (Method details) arising from MFAs within an imaging volume revealed a wide range of activity profiles during locomotion and whisking (Figure 1I). These results show that MFA population activity is highly heterogeneous during spontaneous behaviour.

### Opposing mossy fibre axon responses underlie heterogeneous population activity

Spontaneous behaviour consisted of periods of quiet wakefulness (QW), when the mice rested on the wheel and exhibited little movement or whisking, and bouts of whisking and locomotion, which we defined as the ‘active state’ (AS), which likely encompassed other unobserved behaviours. To investigate the properties of MFA population activity, we sorted activity according to the axonal response during the AS (Figure 2A, Method details). Some MFAs became more active during bouts of locomotion and whisking than during QW (positively modulated, PM), while others were most active during the QW state and became less active during the AS (negatively modulated, NM), while the remainder were not significantly modulated (NSM; Figure 2A,B and S3). This suggests that heterogeneity in the population response properties of MFAs arise, at least in part, from different subsets of axons being active during the QW state and AS. Pronounced heterogeneity in MFA responses across the population was also evident in the temporal domain, with large variations in the latency and duration of the responses of MFAs to locomotion onset, and the activity time courses of some axons lasting longer than the duration of the AS (Figure 2B and S4). The heterogeneity in MFA responses was not due to noise, since axon responses to locomotion onset were reproducible within the same recording (S4). Moreover, the use of real-time 3D movement correction ^51^ in all the recordings confirmed that tissue movement did not underlie MFA response heterogeneity.

**Figure 2.**
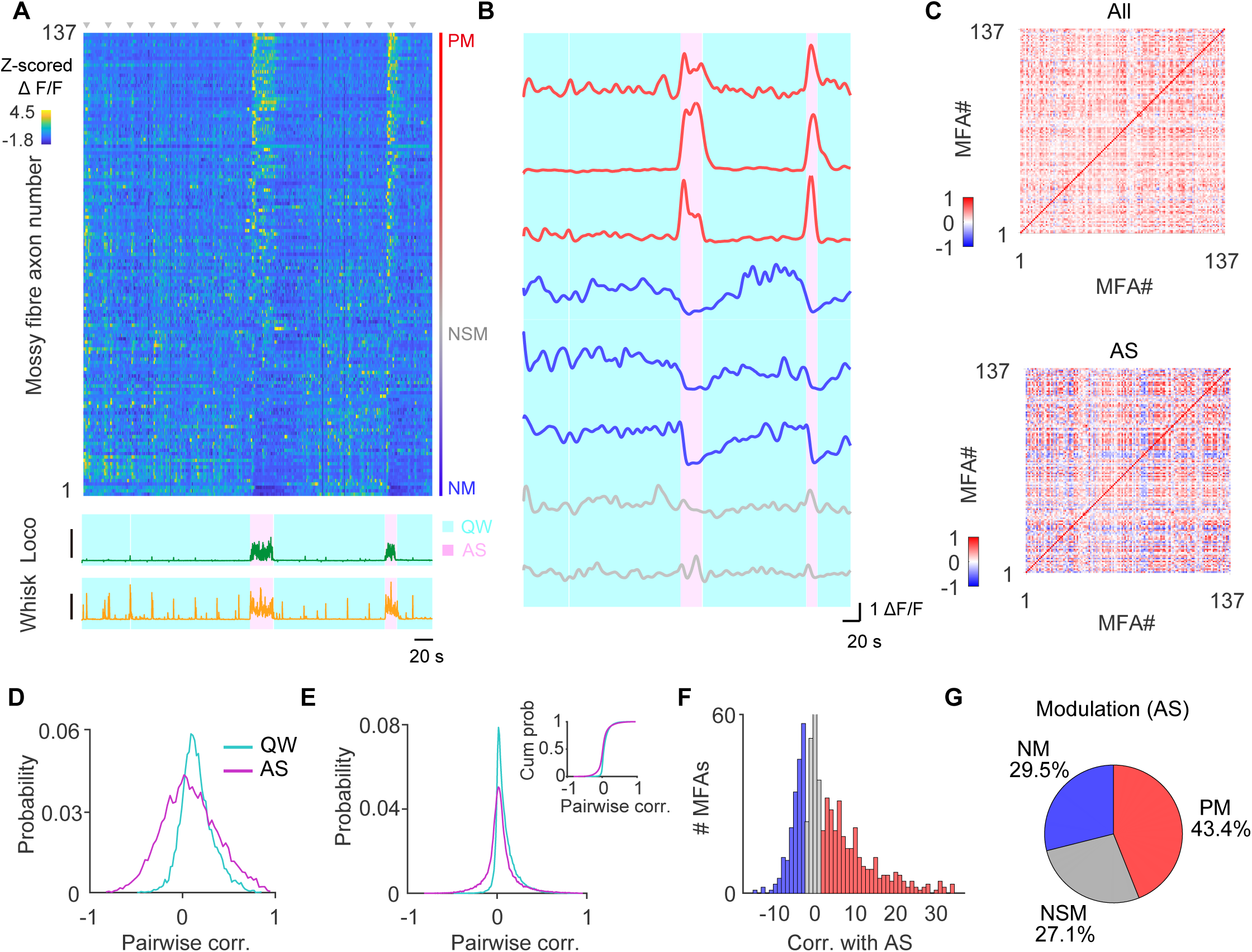
**Bidirectional activity across mossy fibre axons during spontaneous behaviours** (A) Activity of a local population of basal pontine nucleus (BPN) mossy fibre axons (MFAs) during locomotion (Loco) and whisking (Whisk) during a recording session (Z-score of ΔF/F traces (range −1.8 to 4.5, 99th percentile of amplitude distribution). Axonal activity was sorted according to their positive modulation (PM), negative modulation (NM), and non-significant modulation (NSM) during active periods of locomotion and whisking (active state, AS) and quiet wakeful (QW) state. Bottom: The treadmill motion index (Loco) and whisker motion index (Whisk). Scale bar 0.5 cm/s and 0.2 cm/s, respectively. (B) Examples of PM (red), NM (blue) and NSM (grey) BPN MFA responses. (C) Pairwise correlations of MFA activity for the entire session (top) and during the AS (bottom) for the example session shown in (A). (D) Distribution of MFA pairwise correlations for AS and QW states for the example in (A). (E) Distribution of pairwise correlations during QW and AS for all recordings (n = 16, N = 4). Inset shows the cumulative distribution. (F) Normalised bootstrapped correlations for activity of BPN-MFAs exhibiting significant PM (red), NM (blue) and NSM (grey) for the AS. (G) Fractions of significant PM, NM and NSM across animals (n = 16, N = 4).

To further examine MFA reliability and to quantify the temporal response properties across the population, we applied mild and brief (50 ms) air puffs to the whiskers. These evoked reliable responses in a large fraction of the population (S5). Air puff-evoked MFA responses exhibited a wide range of latencies in onset and peak responses across the population. Some MFA responses were rapid (time to peak < 369 ± 11 ms, n = 197 MFAs, *N* = 4), but others exhibited long latencies, reaching a peak more than one second after the air puff (S5). While rapid air puff-evoked MFA responses are likely to reflect the response to this brief sensory stimulus, later components could include startle responses and other air puff-evoked movements. Averaging across the population, the mean time to peak was approximately 1s and the coefficient of variation was 0.5. These results show that BPN-MFAs within local regions of Crus I/II are reliable and exhibit highly heterogeneous temporal responses following a mild air puff to the whiskers.

To further investigate the heterogeneous response properties of MFAs, we examined their pairwise activity. Pearson pairwise correlation coefficient (𝐶𝐶*_p_* ) of axon activity throughout a recording session revealed both positive and negative values (Figure 2C) that became more pronounced during the AS, broadening the distribution of correlations when compared to the QW state (Figure 2D). This change in the shape of the 𝐶𝐶*_p_* distribution during the AS was also evident when data were pooled for all recordings (Figure 2E; KS test, p < 0.001; *n* = 16 sessions, *N* = 4). This suggests that the activity of some axons were positively correlated while others were anti-correlated, and this became more pronounced during the AS.

To relate the responses of MFAs more directly to behaviour, we examined the bootstrapped correlation between MFA activity and behavioural state (QW and AS; Figure 2F). The distribution of correlations between MFA activity and AS had both positive and negative components, consistent with our observations of PM, NM and NSM activity (Figure 2 A,B). To quantify the proportions of MFAs that were significantly PM and NM with the AS, we temporally shuffled the data to determine the level of noise and identified axons with correlations >2SD of the noise (Figure 2G, Method details). Overall, 43% of MFAs were significantly PM and 30% were significantly NM, with the remainder NSM (Figure 2G) during the AS. While the proportions of PM, NM and NSM varied across recordings from Crus I/II, they were present in all animals (S3; *n* = 16, *N* = 4). These results show that a substantial fraction of MFAs are active during both the QW and AS. Moreover, it demonstrates that different behavioural states are represented by different subgroups of MFAs with opposing response properties. This suggests that MFAs entering the cerebellar input layer from the BPN form a widespread bidirectional population code with a substantial fraction of MFAs remaining active even when the animal is behaviourally inactive.

### Geometrical structure of population activity

To gain further insight into the population level properties of MFA activity, we next examined the geometrical structure of activity space during the AS and QW state. Each dimension of the population activity space represents the activity of a different MFA, and each point in space corresponds to a unique pattern of activity across the population. To best capture the overall structure of the population activity subspace, we analysed those recordings with more than 100 MFAs and concatenated sequences of trials to increase the recording duration (Figure 3A). To quantify activity subspace, we decomposed the correlation structure of the population activity into different population modes using Principal Component Analysis (PCA). Visualising the first three principal components (PCs, or modes) of the population activity revealed that QW and AS were associated with distinct subregions of the activity space (manifolds; Figure 3B). Spatially distinct manifolds for AS and QW were evident in all animals and were unaltered by the presence or absence of mild air puffs to the whiskers (S6-S7 and Video S1-S4). Comparison of intra-manifold Euclidean distances suggests that the extent of AS and QW activity space was comparable (*P* = 0.79). However, the distance between the AS and QW manifolds were significantly higher (16%, AS and AS-QW; 19%, QW and AS-QW) than the distance within manifolds (AS and AS-QW, *P* = 6. 1×10^−5^; QW and AS-QW, *P* = 6. 1×10^-4^; Wilcoxon signed-rank test, *n* = 16, *N* = 4; Figure 3C), indicating that the manifolds were spatially separated.

**Figure 3.**
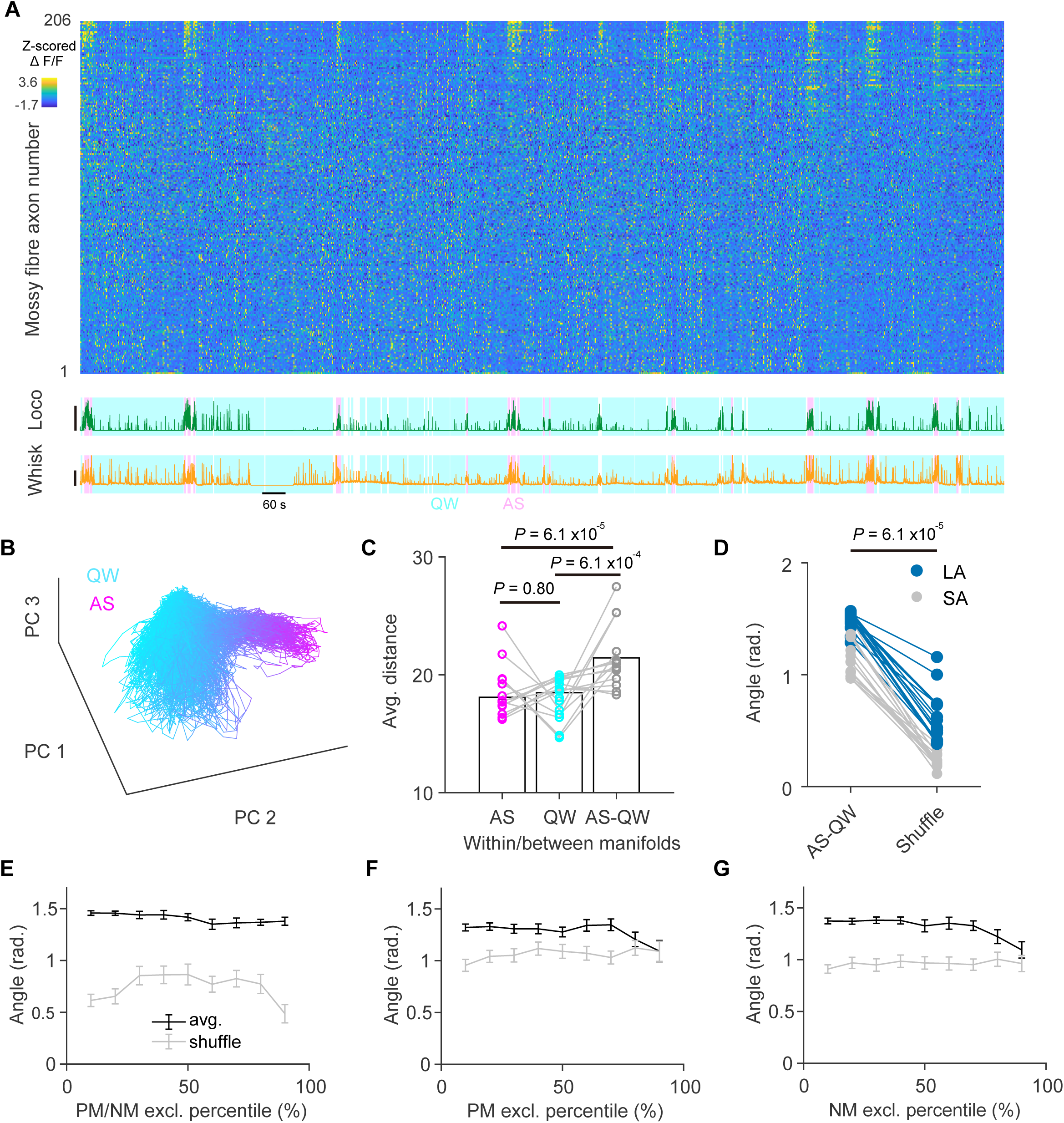
**Geometrical structure of axonal population activity during quiet wakefulness and active behaviours** (A) Extended recordings of mossy fibre population activity (z-scored) during 5 concatenated recording sessions (total duration 2412 s) Bottom: The treadmill motion index (Loco) and whisker motion index (Whisk). Axonal activity was sorted according to the response during active periods of locomotion and whisking (active state, AS) and quiet wakeful (QW) state. Scale bar 0.5 cm/s and 0.1 cm/s, respectively. (B) The first three principal component (PC) trajectories of mossy fiber population activity for the recordings in panel (A). Activity space during quiet wakeful (QW, cyan) and active state (AS, magenta), and the transitions between them. (C) The average Euclidean distance between all pairs of neural activity patterns during different behaviour states within the AS manifold (AS-AS), within the QW manifold (QW-QW), or between the two manifolds (AS-QW). Each circle represents a different experiment (n = 13, N = 4). (D) Angle between QW and AS manifolds in the same experiment together with the null distribution, obtained by calculating the angle between two halves of the data, after temporally shuffling (t = 10s) each experiment (largest angle, LA; smallest angle, SA). (E) Angle between AS and QW manifolds (black data points and line) compared to the mean angle between random halves of the data after shuffling time points (shuffle, grey) as an increasing fraction of NM and PM MFAs are excluded (n = 13, N = 4). (F) Angle between AS and QW manifolds (black) as increasing fractions of the most strongly PM MFAs are excluded. (G) Same as for panel (F) but NM MFs are excluded. Error bars indicate s.e.m..

To quantify how the neural activity subspaces were orientated, we calculated the maximum and minimum angles between the first two PCs of the AS and QW manifolds and compared these values to shuffled controls for each experiment ^24^. Shuffling population activity across time gave a null distribution of angles for each experiment, which was then compared with the angle observed, thereby controlling for measurement noise. The largest angle between the QW and AS activity subspaces was 1.48 ± 0.03 (*P* = 4.9 x 10^−5^; Wilcoxon signed-rank test) radians (Figure 3D), suggesting that these manifold structures were close to orthogonal. Such orthogonal arrangements of activity space are thought to limit interference between the neural representations of different behaviours^52,53,54,55^. We also examined the smallest angle, as this is likely to determine how effectively downstream decoders can distinguish QW and AS manifolds. This was slightly smaller, but still significantly different from the shuffled control (1.18 ± 0.05 radians, *P* = 4.9 x 10^−5^). To test whether subspace orthogonality arose simply from NM and PM MFAs being active during the different behavioural states, we removed increasing fractions of the most strongly modulated axons and recalculated the angle between AS and QW subspaces for each session^24,25^. Surprisingly, as more PM and NM MFAs were excluded, the angle between these subspaces remained relatively constant and significantly larger than in the shuffle control (*P* < for 0th through 90th percentile of PM and NM MFAs excluded, Wilcoxon signed-rank test with Bonferroni correction for multiple comparisons; Figure 3E). Furthermore, the near-orthogonal geometry remained when a substantial proportion of either PM or NM MFAs were excluded (Figure 3F and Figure 3G, respectively). These results suggest that the orthogonal arrangement of the subspaces does not entirely arise from different subsets of MFAs with opposing response properties, but that PM and NM axons contribute to both AS and QW manifolds.

### Contribution of mossy fibre axons to population activity modes

To investigate the correlation structure of MFA activity in more detail, we examined the amplitude (eigenvalues) of the different population activity modes. Plotting the first 20 PCs revealed an eigenspectrum, with the amplitudes of the eigenvalues decreasing relatively gradually (Figure 4A). Indeed, the top mode had an amplitude only 1.4 - fold larger than the next mode (PC2/PC1 = 0.71 ± 0.034, *n* = 13, *N* = 4). This suggests that the MFA population activity consists of a large number of independent modes.

**Figure 4.**
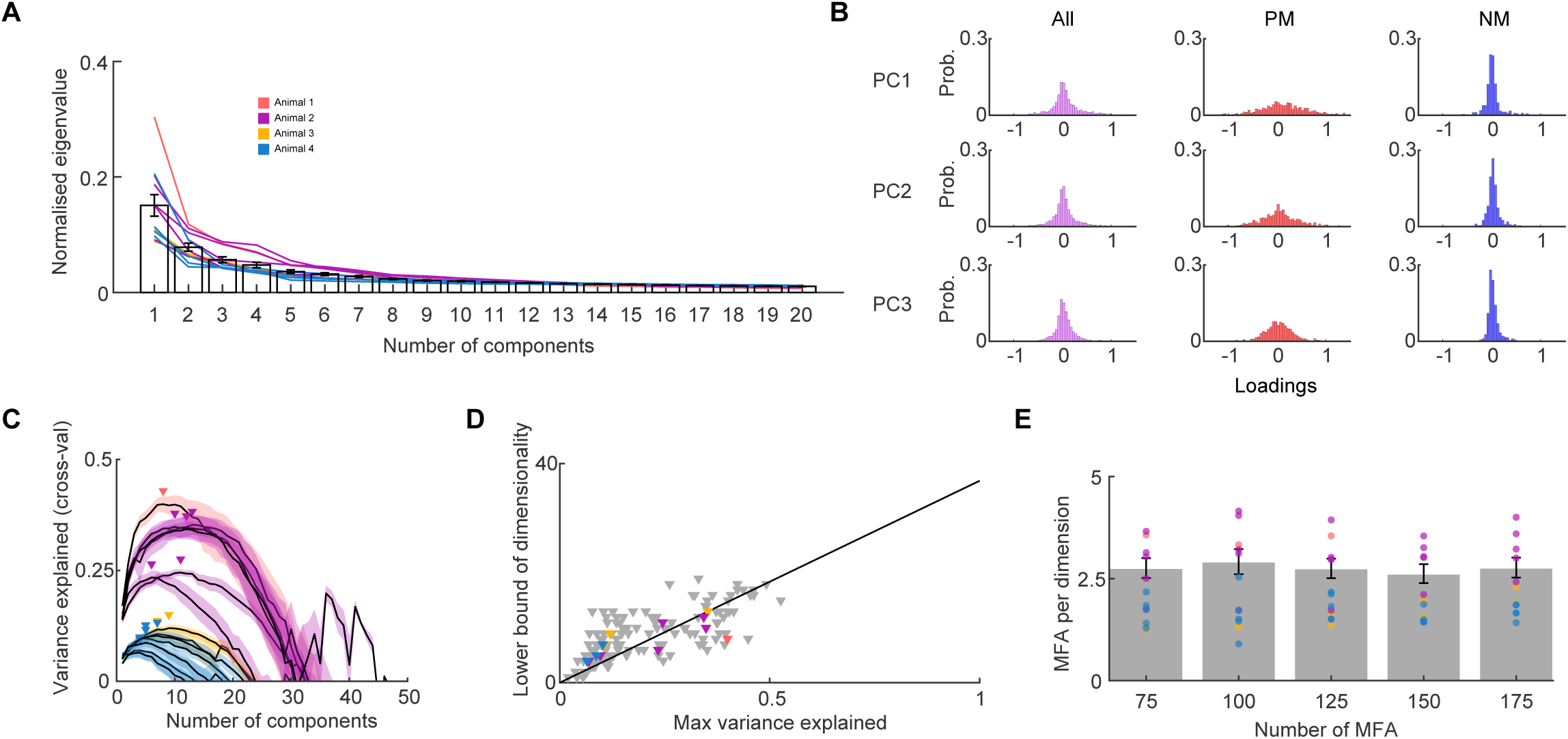
**Dimensionality of mossy fibre population activity during spontaneous behaviours** (A) Normalised eigenspectrum showing the first 20 principal components (PCs, modes) across all recording sessions for z-scored ΔF/F traces across all sessions (n = 13, N = 4, each colour represents a different animal). (B) Distribution of loadings for the first three PCs for all, PM and NM MFAs for z-scored ΔF/F activity during active states. (C) Relationship between cross-validated explained variance and PCs for all sessions for z-scored ΔF/F activity (n = 13, N = 4). Colours indicate different animals, arrows indicate peak explained variance for each session. (D) Relationship between the lower bound of the dimensionality and the maximum variance explained (Vmax). Grey and coloured arrowheads indicate individual subsamples of held-out data and means for each experiment (peaks in c), respectively. Linear extrapolation predicts that 38 dimensions are necessary to explain all of the variance for populations of 75 mossy fibres. (E) The ratio of number of MFAs to the extrapolated dimensionality for different subsampled population sizes: 75 MFAs (n = 13, N = 4) to 175 MFAs (n = 12, N = 3).

To examine the contribution of MFAs to the population modes, we calculated their loadings, since they reflect the weighted linear combination of the individual MFA activity. The fact that loadings were positively and negatively distributed around zero for each PC (Figure 4B) suggests that both positive and negatively correlated MFAs contribute to the top 3 population modes, consistent with our geometric analysis of the AS and QW manifolds. To determine the linear embedding dimensionality^56^ of the MFA population code, we determined the number of population modes required to account for the maximal explained variance during spontaneous behaviours for recordings with ≥75 MFAs, using cross-validated PCA^57,104^.

This provides an estimate of the dimensionality of the population activity that is shared across MFAs (shared dimensionality). The relationship between cross-validated explained variance (CVEV) and number of population activity modes for each session increased to a peak, before declining due to noise or other non-shared variability (Figure 4C). Although the extent of variance explained varied across sessions and animals, the location of the peak, which provides a lower bound estimate of the shared dimensionality^24^, tended to increase with the CVEV.

Linear extrapolation of the relationship between the lower bound of the dimensionality and variance explained, suggested that 38 dimensions are required to explain the full variance for a random sub-sampled population of 100 MFAs during spontaneous behaviours (Figure 4D). An alternative assay of embedding dimensionality of the activity subspace that includes independent MFA activity and noise, can be estimated from the total variability after adjusting for relative amplitudes of the population modes^103^. This effective dimensionality measure was also high for the same group of MFAs (78 ± 6.2; *n* = 13, *N* = 4). Since measures of dimensionality depend on the numbers of neurons recorded, we used the extrapolated lower bound estimate of the shared dimensionality to calculate the number of dimensions per MFA^24^.

This gave a surprisingly low value of 2.8 MFAs per embedding dimension for randomly selected subset populations containing between 75 and 175 axons across sessions and animals (Figure 4E; or 3.2 MFBs per dimension for 125 to 275 MFBs). The inverse of this measure is perhaps more intuitive, in that it provides an estimate of the fraction of the activity state space that is utilised (linear embedding dimension/total number of MFA = 0.31). These results show that the embedding dimensionality of MFA activity is high during spontaneous behaviours and that few MFAs encode each dimension. This suggests that the ponto-cerebellar projection has a high capacity to encode different behaviours and states.

### Decoding behavioural variables from population activity

The high embedding dimensionality of the MFA population activity is likely to reflect a wide range of encoded variables, including observed and unobserved behaviours, as well as internal states and computations. To understand how MFA population activity is related to the observed behaviours, we decoded whisking, locomotion and behavioural state from behaviourally active sessions using population activity modes as predictors. To ensure robustness against co-linearity and overfitting, we used cross-validated ridge regression to examine the relationships between whisking, locomotion, behavioural state and population modes (Figure 5A). Across animals and sessions, the top PC was weakly correlated with locomotion, whisking and the behavioural state accounting for only 0.19 ± 0.06, 0.31 ± 0.10 and 0.28 ± 0.09, of the behavioural CVEV, respectively (*n* = 13, *N* = 4). This suggests that the largest population activity mode was not always dominated by the observed behaviours or AS. Adding more population modes as predictors in the regression increased the CVEV of the behaviours (Figure 5A). Plotting the CVEV as a function of the number of modes revealed a saturating function with a shallow slope above 50 MFAs. We therefore used a stopping function to estimate the ‘optimal’ number of population modes (Method details) for each behavioural variable (Figure 5B,C). Decoding using this optimal number of PCs as predictors led to substantially better decoding performance than for the first PC for locomotion (*P* = 2.4 x10^-4^), whisking (*P* = 2.4 x10^-4^) and AS (*P* = 4.9 x10^-4^; (Wilcoxon signed-rank test, *n* = 13, *N* = 4; Figure 5D). This improvement was not due to an increased number of parameters because decoding performance was cross-validated. These results suggest that the top population mode is weakly correlated with behaviours and the activity state of the animal and that information on specific behaviours is distributed across multiple population modes.

**Figure 5.**
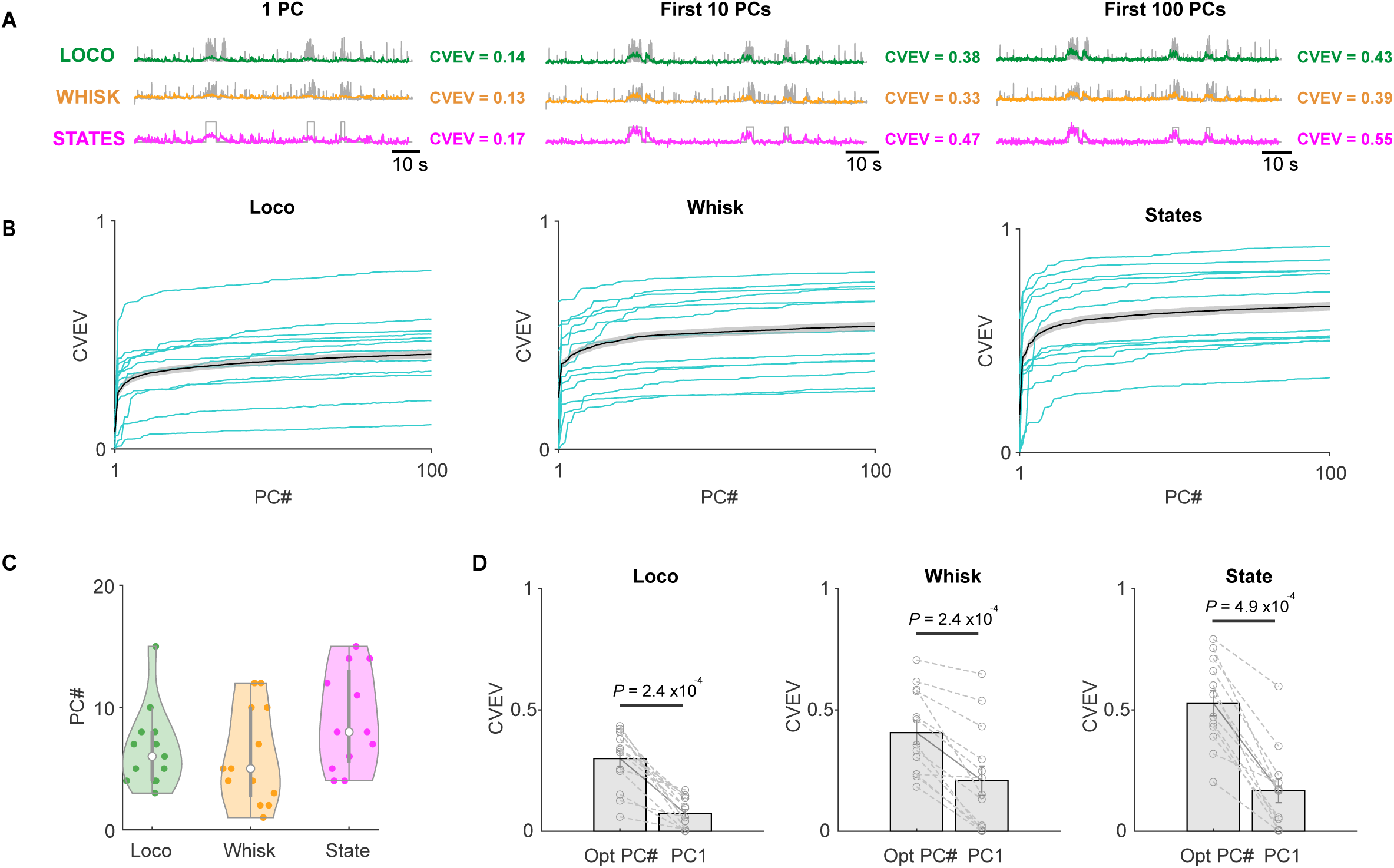
**Relationship between population activity modes and behaviour** (A) Example of using different numbers of principal component population activity modes to predict locomotion (Loco), whisking (Whisk) and active state (AS) using cross-validated ridge regression. Cross-validated explained variance (CVEV) indicated on the right of traces. (B) Relationship between CVEV and population modes for individual sessions with significant activity for regressed behaviour (cyan) and overall mean (black; Loco and AS n = 12, Whisk n = 13, N = 4). (C) Estimate of optimal number of modes for ridge regression for different behavioural variables across all sessions (n = 13, N = 4). (D) Comparison of CVEV for locomotion, whisking and AS when using PC1 and the optimal number of modes as regressors. (P values Wilcoxon signed-rank test).

### Some axons are highly behaviourally informative

To investigate whether some MFAs are more behaviourally informative than others, we decoded behavioural state, whisking and locomotion with ridge regression using the activity (Z-score) of MFAs as predictors. The MFAs with the highest beta coefficients were highly behaviourally informative, accounting for a large fraction of the CVEV for locomotion, whisking and AS (Figure 6A). On average, the most behaviourally informative MFA in each session for locomotion, whisking and behavioural state accounted for 0.22, 0.26 and 0.45 of the CVEV, respectively (*n* = 13 sessions, *N* = 4 animals). Interestingly, these MFAs were all PM. Nevertheless, decoding performance increased with the number of MFAs included as predictors, and could account for a large fraction of the CVEVs for behavioural state, locomotion and whisking with a hundred or so MFAs (Figure 6B). To quantify the minimal number of axons necessary for optimal decoding, we next applied lasso regression (L1 regularisation; Method details). This gave a minimum number of MFAs of 57 ± 10, 41 ± 8, 146 ± 6 for locomotion, whisking and behavioural state across sessions (Figure 6C). The CVEV of these was significantly higher than for the most informative MFA in each session (*P* = 2.4 x10^-4^, 2.4 x10^-4^, 4.9 x 10^-4^, Wilcoxon signed-rank test, *n* = 13, *N* = 4). These results show that while behavioural information is distributed across multiple MFAs, some individual MFAs are highly behaviourally informative.

**Figure 6.**
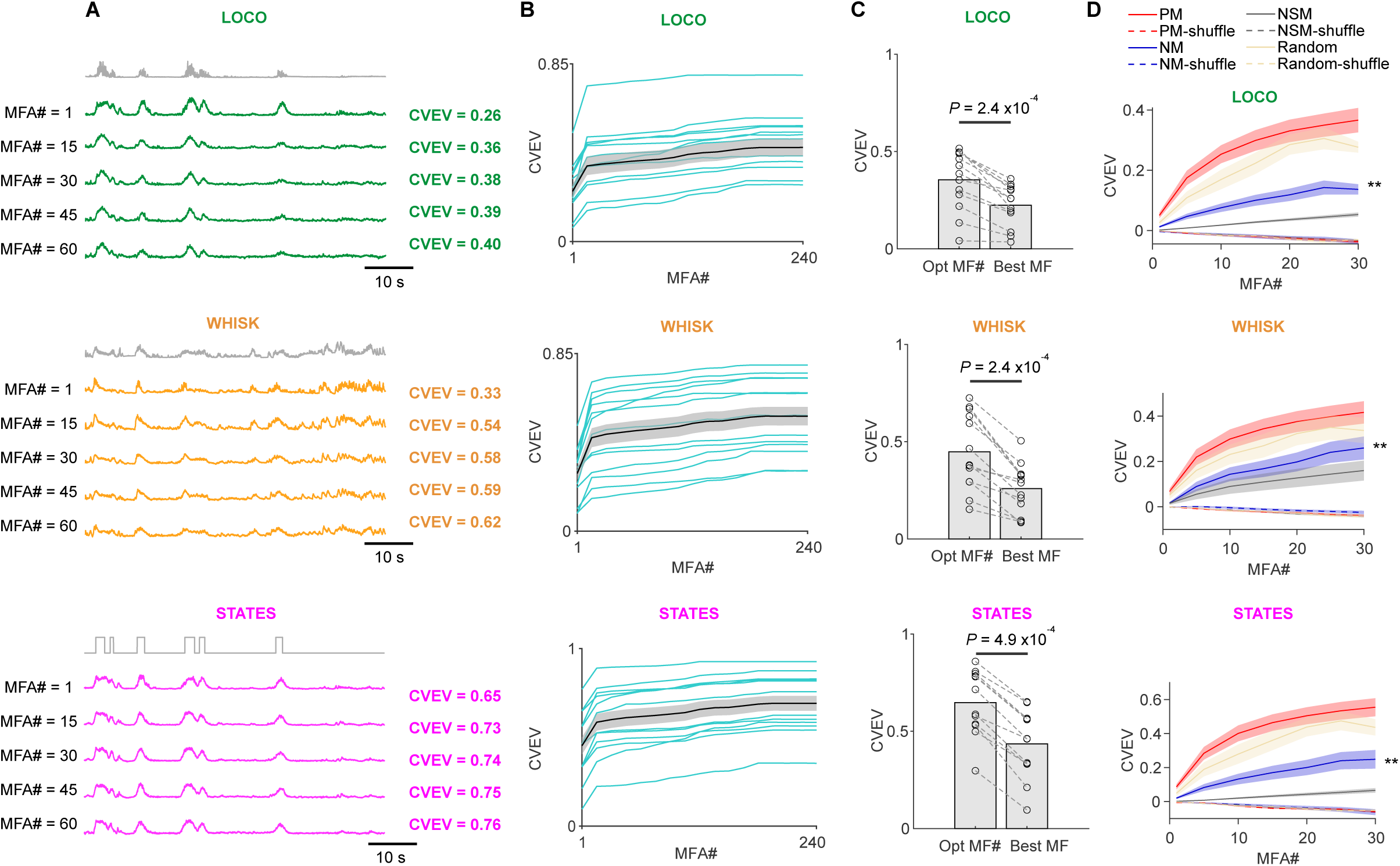
**Relationship between mossy fibre activity and behaviour** (A) Examples of using different numbers of the most informative mossy fibre axons (MFAs) to predict locomotion (Loco), whisking (Whisk) and active state (AS) using cross-validated ridge regression. Cross-validated explained variance (CVEV) for an individual session, indicated on the right of traces. (B) Relationship between CVEV and number of MFAs for sessions with significant activity for regressed behaviour for individual sessions (cyan) and overall mean (black; Loco and AS n = 12, Whisk n = 13, N = 4). (C) Comparison of CVEV for locomotion, whisking and AS for the minimal number of MFA regressors identified with cross-validated lasso regression and the most informative single MFA (P values Wilcoxon signed-rank test). (D) Relationship between CVEV and number of randomly selected MFAs for locomotion (top), whisking (middle) and state (bottom) from either the whole MFA population (Random) or the PM, NM and NSM subgroups using cross-validated ridge regression, together with their temporarily shuffled controls (shuffle). Statistically significant with respect to shuffled controls (Wilcoxon signed rank test, P values 0.0078 at 30 MFAs, ** P <0.01).

### Positively and negatively modulated mossy fibre activity is behaviourally informative

To investigate how PM, NM and NSM MFAs contribute to behavioural representations, we performed ridge regression on 1 to 30 axons randomly subsampled from these groups and compared their decoding performance to a randomly subsampled group from all axons. PM MFAs were consistently more informative than NM MFAs alone, NSM MFAs or shuffled controls for the measured behaviours (Figure 6D; but could be more informative about other parameters such as pose). Nevertheless, the CVEV of NM MFAs was significantly higher than the shuffled control for locomotion (*P* = 0.0078), whisking (*P* = 0.0078) and behavioural state (*P* = 0.0078) (Wilcoxon signed-rank test, *n* = 13, *N* = 4, 30 NM; Figure 6D), indicating that NM MFAs convey behavioural information. Random mixtures of PM, NM and NSM MFAs were substantially more predictive than NM or NSM MFAs alone (for 30 MFA, CVEV 27.6% for locomotion and 33.7% for whisking and 43.6% for behavioural states; Figure 6D). These results suggest that both PM and NM MFA populations convey behavioural information, with PM MFAs being more informative on average, consistent with the most behaviourally informative single MFAs being PM (Figure 6D). These results show that behavioural information is distributed across MFAs with opposing response characteristics forming a bidirectional population code.

### Spatial properties of mossy fibre activity

To better understand how synaptic input from PM, NM and NSM axons could be integrated by downstream GrCs during behaviour, we examined their spatial properties, since spatial clusters of similarly tuned MFBs could lead to more reliable and homogeneous GrC activation^58^. For individual GrCs, the relevant spatial scale for MFA synaptic inputs is the maximal span of their dendritic tree, which is approximately 40 μm^22^. We analysed synaptic boutons rather than axons as their location can be described by points. Visualisation of the location of PM, NM and NSM MFBs within an imaging volume suggests they are spatially mixed in 3D space (Figure 7A). To quantify whether PM and NM MFBs were differentially distributed through the imaging volume, we compared their distributions in X, Y, and Z dimensions (Figure 7B). However, there was no significant difference between these distributions (*P* = 0.95, 0.24, 0.07; KS test). To further investigate whether MFBs were spatially clustered, we compared their 3D spatial distribution to a completely random distribution of points using Ripley’s L function^59,105^. For each MFB type L increased with distance between MFBs, remaining close to the unitary slope line up to a radius of 50 μm, as expected for a completely spatially random distribution (Figure 7C; *n* = 13, *N* = 4). We next investigated whether MFB activity is spatially clustered during spontaneous behaviours by examining the 𝐶𝐶*_p_* of MFB activity. This revealed a uniform spatial structure on the 200 μm scale, both in the QW state and AS (Figure 7D; *n* = 13, *N* = 4). When PM, NM and NSM MFBs were examined separately, they also had a spatially invariant 𝐶𝐶_0_ structure during behaviours (Figure 7E). These results suggest that PM, NM and NSM MFBs are intermingled on spatial scales that are relevant for integration by individual GrC cells and that there is no significant spatial dependence in the correlation structure of their activity.

**Figure 7.**
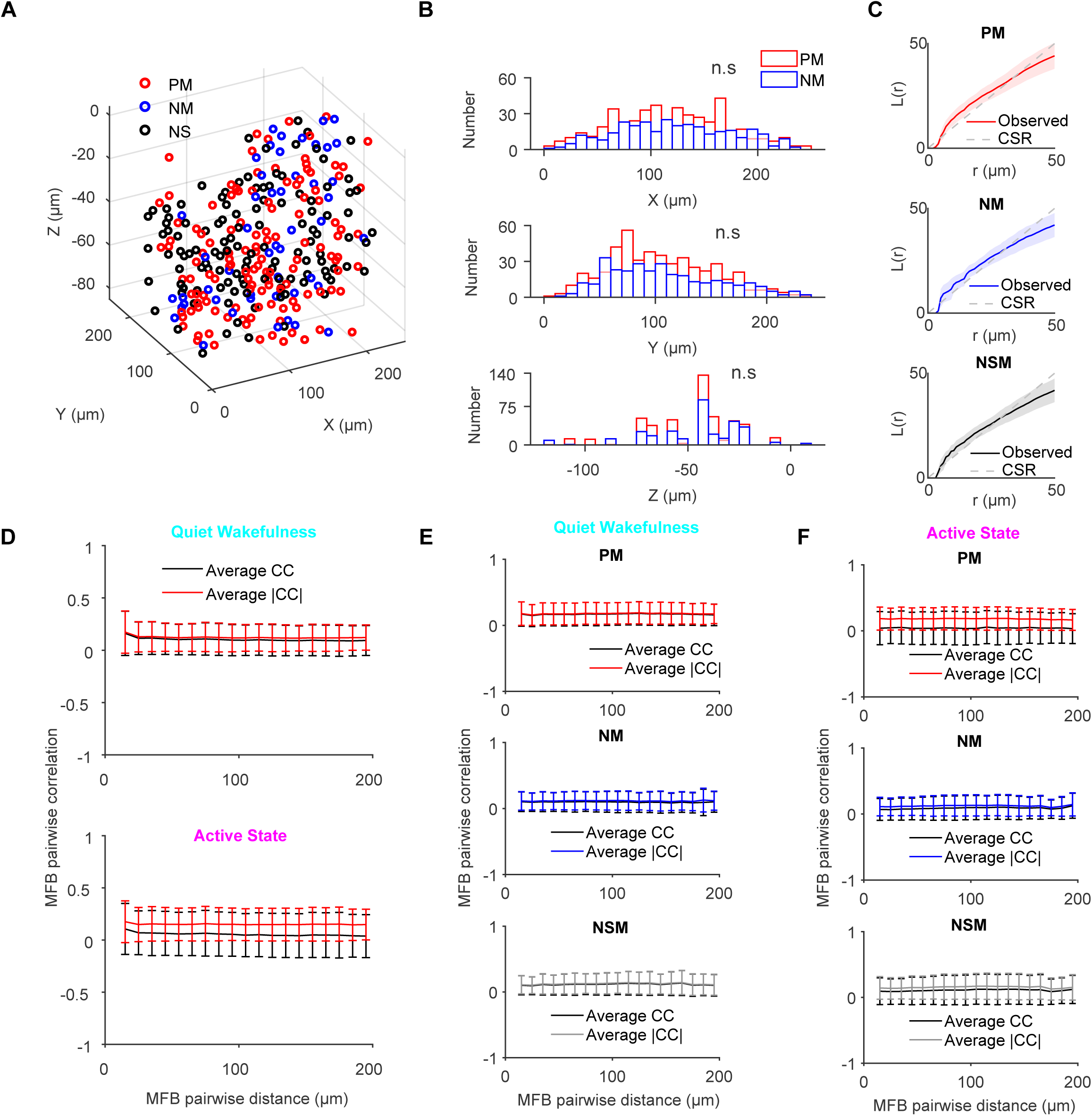
**Spatial properties of mossy fibre boutons in the granule cell layer** (A) Locations of positive (PM, red), negative (NM, blue) and not significantly (NSM, grey) modulated mossy fibre boutons (MFBs) in an individual imaging volume. (B) Spatial distribution of all PM and NM MFBs across all animals, in X (top), Y (middle) and Z (bottom) dimensions (P values KS test, n = 16, N = 4). (C) Ripley’s L function (L(r)) for PM (top), NM (middle), and NSM (bottom) MFBs across different radii (r). The shaded regions represent the s.e.m calculated across animals (N = 4). The expected distribution assuming Complete Spatial Randomness (CSR, dashed line). (D) Average pairwise cross correlation (CC) of MFB activity as a function of their distance, during the QW (top) and AS (bottom) for both values and magnitudes. (E) Distance dependence of pairwise CC for PM (top), NM (middle), and NSM (bottom) MFB activity during quiet wakefulness. (F) Same as for (E), but for active state.

### Implications for synaptic integration in granule cells

The intermingling of PM, NM and NSM MFBs on short spatial scales and the lack of spatial structure in MFB activity in the cerebellar input layer suggest that GrCs could be innervated by different combinations of PM, NM and NSM axons. As it is currently technically challenging to test this experimentally (i.e. to record the activity of all the presynaptic MFAs that innervate an individual GrC, together with its postsynaptic activity) we developed a simple model of MFB-GrC connectivity using the measured locations of PM, NM and NSM MFBs and by placing 300 GrCs randomly within the imaging volume. Connections were then made between each GrC and its 4 closest MFBs. This process was repeated ten times to determine the frequencies of occurrence of different combinations (Figure 8A; Method details). The largest proportion of GrCs were innervated by 2 PM, 1 NM, and 1 NSM MFAs (13.3%), while the next largest proportions were GrCs with combinations of 2 PM and 2 NSM MFAs (11.5%). GrCs with purely PM MFAs and purely NM MFAs made up 6.5% and 3.2%, respectively, whilst the proportion of GrCs that were purely innervated by NSM MFAs was 1.4%. These proportions are qualitatively similar to theoretical predictions for 4 random samples of 3 object classes without replacement (Figure 8; Method details), consistent with MFBs being randomly distributed.

**Figure 8.**
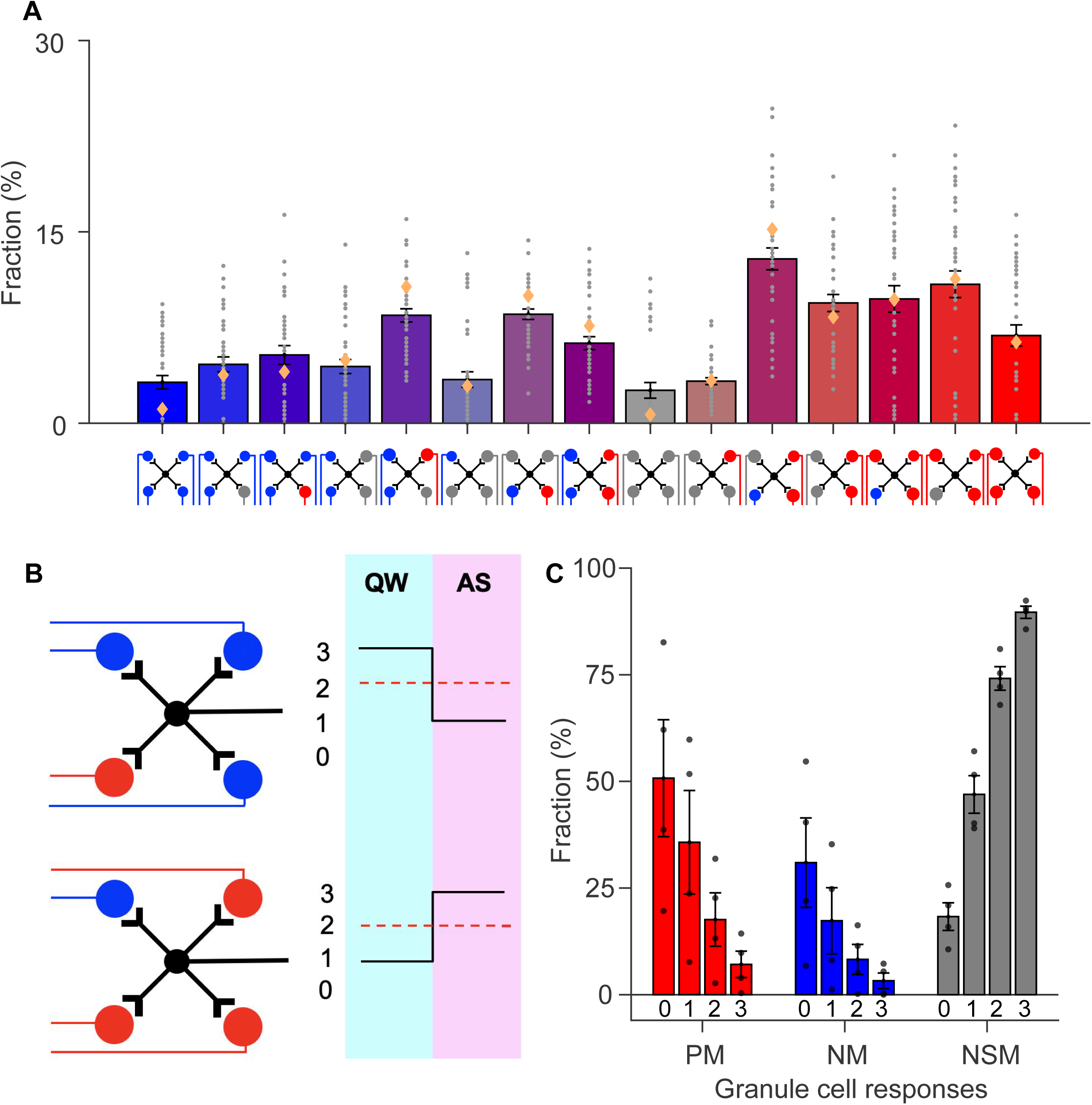
**Simple model of mossy fibre to granule cell connectivity** (A) Predicted proportions of GrCs innervated by different combinations of PM, NM and NSM MFAs using measured locations of MFBs and assuming nearest neighbor connectivity. Bars indicate the average over 10 model instances per animal (4 animals total), with each instance using random placements of 300 GrCs. Orange diamond dots indicate theoretical value for each combination assuming random sampling of 4 MFAs from 3 functional types, without replacement. (B) Schematic illustration of NM GrC with 3 NM (blue) and 1 PM (red) MFAs (top, left) and activity during QW and AS (top, right) assuming synaptic weights of 1 and 0 during different states. Dashed line indicates GrC spike threshold (Th) of 1. Bottom: the same, but for PM GrC with 3 PM and 1 NM MFAs. (C) Predicted proportions of GrCs with different response properties (PM, NM, NSM) for different GrC thresholds (Th). For GrCs to be PM or NM their activity must change between the QW and AS and their inputs sum to a value > Th in one or both of the states (the proportions for GrC for Th = 0, PM 52.0%, NM 30.7%, NSM 17.4%. For Th = 1, PM 33.6%, NM 18.2%, NSM 48.3%. For Th = 2, PM 17.5%, NM 8.2%, NSM 74.3%. For Th = 3, PM 6.5%, NM 3.2%, NSM 90.3%.

To relate this expected MFA-GrC connectivity to GrC activity, we made the simplifying assumption that synapses made by PM, NM and NSM MFAs had an amplitude of 1 or 0. NM MFAs were active (1) during the QW state and inactive (0) during the AS. PM MFAs were active during the AS and inactive during the QW state, while NSM MFAs were always inactive. For each of the combinations, we then quantified the number of NM MFAs that were active during QW and the number of PM MFAs that were active during the AS (Figure 8B). If the number of active MFAs per GrC increased between the QW and AS, the GrC was positively modulated; and if it decreased, it was negatively modulated. For simple linear integration, GrCs with PM and NSM MFA input combinations, and those with more PM than NM inputs, will generate positively modulated GrC responses during transitions from the QW to AS (53%). GrCs with NS and NSM MFA input combinations, and those with more NM than PM inputs, will generate negatively modulated GrC responses (31%), while GrCs with NSM only or equal PM and NM input combinations will generate GrCs with NSM GrC responses (16%). These results should reflect the linear subthreshold GrC voltage response properties. We then investigated how innervation by different MFA combinations might influence GrC firing. A threshold of zero, where only one active MFA is required during the QW or AS, gave a similar proportion of PM, NM and NSM GrCs as the linear subthreshold case, due to the binary nature of the inputs (Figure 8C). However, increasing the GrC spike threshold increased the proportion of inactive GrCs at the expense of PM and NM GrCs (Figure 8C), with inactive fractions reaching 18%, 47%, 74%, and 90% when the threshold was set to 0, 1, 2, and 3, respectively (Figure 8C; assuming NSM MFAs are inactive rather than continuously active). However, the NM/PM ratio remained relatively constant (0.47 - 0.59). For a threshold of 3, only GrCs innervated with purely PM MFAs or purely NM MFAs responded, with all other combinations remaining silent. Although these predictions rely on simplifying assumptions, and are qualitative, they show that intermingled MFAs with opposing response properties are expected to generate bidirectional responses in the downstream GrC population, with a consistently larger proportion of PM than NM responses, as recently observed during spontaneous behaviours^24^.

## Discussion

### Summary

Our experiments provide the first measurements of population activity of mossy fibre input to cerebellar cortex. MFAs originating from the BPN exhibited highly heterogeneous responses that were either positively or negatively modulated during locomotion and whisking. Analysis of the axonal activity space revealed high dimensional representations and orthogonally arranged manifolds during active and quiet wakeful states. A surprisingly large fraction of the population activity state space was utilised during spontaneous behaviour, and some MFAs in the population were highly behaviourally informative. Mossy fibre boutons with opposite signed activity were intermingled in the granule cell layer and there was no detectable spatial correlation in their activity over hundreds of micrometres. A simple model of GrC integration suggests that the bidirectional MFA code is transmitted effectively to GrCs, which sample different combinations of MFAs with opposing response properties. Our results establish the basic functional properties of pontine mossy fibre axon population activity and suggest that the cortico-pontine input to the cerebellar cortex utilises a high capacity, low redundancy, bidirectional code to convey sensorimotor information during spontaneous behaviours.

### Relationship between BPN-MFA population activity and individual mossy fibre response properties

Our recordings from hundreds of MFAs during behaviours expand our view from single MFAs to the population level, revealing properties that could not be inferred from previous recordings of individual MFAs^40^. Our results show that behavioural information is distributed across BPN-MFAs with opposite signed response properties and that their activity is surprisingly varied, having a high embedding dimensionality and broad latency distribution. Moreover, the population activity occupies distinct, orthogonally arranged, activity subspaces during the QW and AS. Nevertheless, some BPN-MFAs are highly informative about specific behavioural variables, such as whisking and locomotion (e.g. Figure 6), consistent with electrophysiological recordings from individual MFAs and GrC EPSCs, which showed putative MFAs can be tuned for joint angle^39^, gait during locomotion^40^, angular head velocity^42^ and whisking^43^. Our results establish that during complex behaviours sensorimotor information from the neocortex is conveyed to the cerebellar cortex with a high capacity, low redundancy, bidirectional BPN-MFA population code.

### Properties of BPN mossy fibre axon population activity

Our analysis reveals that MFAs that are positively modulated during active behavioural states form a majority, while those that are more active during the QW state, and are negatively modulated during the AS, form a substantial minority. Cross-validated regression analyses show that both subpopulations are behaviourally informative, but that PM responses are more informative, on average, for the whisking and locomotion behaviours we have measured (but note NM could be more informative about other factors such as pose). Consistent with this, the most informative individual MFA in each circuit for whisking and locomotion was consistently from the positively signed subgroup. These findings establish that BPN-MFAs utilise a bidirectional population code for cortico-ponto-cerebellar signalling. Coding strategies that split information into ON- and OFF-type responses have been reported in other vertebrate and invertebrate circuits, including those involved in visual processing^60,61^, thermosensing^62^, chemosensing^63^, and auditory processing^64^. ON-OFF responses have also been reported in motor cortex^65,66,67^. Theoretical analysis of coding strategies has shown that splitting pathways into ON- and OFF streams can transmit the same amount of information as purely ON systems, but more efficiently, with fewer spikes^68^. This may confer a metabolic advantage as MFAs fire for sustained periods^39,42^ over a wide frequency bandwidth^41,45^ and MFBs are packed with mitochondria^69^. These observations suggest the bidirectional BPN-MFA population code is likely to have evolved, at least in part, to provide an energy efficient solution for transmitting the extensive sensory and motor command signals from the neocortex to the cerebellar cortex, thereby enabling it to be continuously updated on the current and future state of the animal.

An unexpected feature of the BPN-MFA population activity is the high embedding dimensionality of the representation during behaviours, with an average of 3 MFAs encoding each dimension. To set this in context, the theoretical maximum number of dimensions a neural circuit can encode (i.e. the number of degrees of freedom of the neural activity) is equal to the number of neurons. This occurs when the activity of each neuron is independent of all the others, resulting in an eigenspectrum that is flat, with each population mode of equal amplitude. While variability arising from noise can inflate some measured dimensionality^18,70^, our cross-validated approach accounts for noise and provides a lower-limit estimate of the embedding dimensionality^57^. Nevertheless, our estimate is still surprisingly close to the theoretical upper bound indicating that a large fraction of the activity state space is utilised during complex behaviours. Recent recordings from neocortex have revealed that embedding dimensionality scales linearly with the number of neurons^71^. The constant ratio of dimensions per axon across different sample sizes indicates the embedding dimensionality of the population code also scales with BPN-MFA number. These results suggest that BPN-MFAs are relatively decorrelated and that their population code exhibits a low level of redundancy during behaviours.

As our PCA was applied throughout sessions, it is likely that some of the network activity is unrelated to the behaviours we have measured. Indeed, the top population mode was relatively poorly correlated with locomotion, whisking and AS (𝐶𝐶*_p_* = 0.18, 0.30, 0.28 𝐶𝐶 = 0.16, 0.31, 0.26). While our results reveal high dimensional activity during a mixture of behaviours, the BPN-MFA population should also be capable of conveying the lower dimensional, more redundant representations that emerge during skilled learning of simple tasks^38^. Our finding that the major MFA input to the cerebellar cortex can encode a large number of independent variables fits well with theories of cerebellar function that combine a wide range of sensory and motor command signals to generate internal models that predict the state of the animal^1,30^.

### Origins of high dimensional bidirectional population activity

A major component of the BPN-MFA activity we recorded in CrusI/II is likely to have originated from sensory and motor areas of the neocortex, since viral tracing and functional mapping studies show that the BPN receives input from these regions of the neocortex and that MFAs from these neurons project densely to the cerebellar hemispheres^17,9^. Interestingly, even the earliest electrophysiological recordings from the motor cortex noted the heterogeneity in the neuronal response properties and that some cells are activated during behaviours, while others are active during quiet wakeful states^66^. More recent imaging of corticospinal neurons in mouse motor cortex suggests there are larger fractions of quiescent active than movement active neurons, and that indiscriminately active neurons make up the majority^65^, a result consistent with Neuropixels recordings from Layer 5^67^. Interestingly, patch clamp recordings from L5b neurons in the mouse motor cortex revealed bidirectional responses with a similar ratio of suppressed to enhanced responses to the NM/PM BPN-MFAs we observe in the cerebellar cortex^72^ (71% vs 68% respectively). The striking similarity of their functional properties suggests that the bidirectional responses of BPN-MFAs are likely to have a neocortical origin^73^.

There are several other properties of the BPN-MFA population code that likely reflect a neocortical origin. These include the mixed selectivity of pyramidal cells^25,65^, the high embedding dimensionality of the population coding in sensory areas^57^ and across the neocortex^71^, as well as the orthogonal manifold structures observed during active behaviours and quiet wakefulness in motor areas^52,53,74^, which have been proposed to separate neural activity that drives behavioural output from activity that represents internal computations, such as motor preparation^55^. Moreover, high pairwise correlations between L5b pyramidal neurons in motor cortex and cerebellar granule cells suggests strong connections can be formed after learning a skilled forelimb task^38^. While these similarities between neocortical and BPN-MFA population codes might suggest BPN-MFAs simply convey a copy of the neocortical output to the cerebellar cortex, several factors are likely to transform neocortical activity as it flows through the BPN^8,16,15^. These include its characteristic 3D layered structure^12,75^ and electrophysiological properties^76,73^ which are likely to perform decorrelation and mixing. Moreover, the excitatory feedback connections from the cerebellar output nuclei to BPN^14,15^ could extend the timescale over which pontine neurons can sustain their firing, contributing to the large temporal heterogeneity we observe in BPN-MFA responses (S4 and S5). Thus, the spatio-temporal heterogeneity, geometric structure, and high dimensionality of the BPN-MFA population activity are likely to arise from a combination of neocortical input, thresholding by BPN neurons and feedback signals from the cerebellum.

### Implications for cortico-ponto-cerebellar communication

Our results indicate the neocortex communicates with the cerebellum using a low redundancy, high capacity, bidirectional population code. The fact that BPN-MFA activity is high dimensional suggests the cerebellar input layer receives a variety of sensorimotor input from the neocortex, even on the relative local spatial scales of our imaging volume (∼200 μm). The projection from the motor cortex has long been proposed to play a key role in motor control by conveying motor command signals (efference copy/corollary discharge) to the cerebellar cortex^3^. Combining these with proprioceptive and sensory inputs, including those from sensory areas of the neocortex is thought to enable the cerebellum to form internal models, perform sensorimotor prediction^1,2,6^ and rapid motor learning^77^. Recent theoretical work suggests that the convergent-divergent anatomy of the cortico-ponto-cerebellar pathway could facilitate this functionality through structured compression^16^. Our results showing that pontine MFA inputs to the cerebellar cortex exhibit high capacity and low redundancy during complex behaviours is consistent with the proposal that the pontine nuclei optimally compress a more distributed cortical representation of sensorimotor information.

A striking feature of the BPN-MFA population response is the wide range of latencies to mild air puffs (S5). While the fastest responses are likely to reflect sensory input and startle responses, some responses take more than a second to reach peak activity. This is much longer than the 100 ms or so temporal resolution of GCaMP6 imaging and likely reflects slower air puff-evoked behaviours or internal computations. The large latency distribution of responses together with the sustained activity, and pauses in activity, of individual BPN-MFAs could provide an extended temporal basis for learning^4^, that stretches beyond the timescales of MFA-GrC short-term plasticity^27,44,78^. These temporal dynamics could also contribute to the cerebellar learning of intervals on the timescale of 1s and beyond^79^.

### Implications for synaptic integration in the granule cell layer

Our finding that BPN-MFAs with different response properties are locally intermingled in the input layer of CrusI/II, suggests that individual GrC cells can be innervated by different combinations of MFAs with opposing signs. Our simplified model of the cerebellar input layer, which was based on measured anatomical properties, suggests that the bidirectional response properties of the BPN-MFA population activity are transmitted effectively to downstream GrCs, irrespective of the level of their spike threshold, a prediction that matches the measured bidirectional response properties of GrCs during spontaneous behaviours^24^. It may seem surprising that some GrCs are expected to be innervated by opposite signed MFA inputs. However, rather than GrCs simply not changing their activity as a result of opposing MFA activity during QW and AS, opposite signed MFA inputs with different latencies, amplitudes and synaptic dynamics^27^ are likely to contribute to the highly heterogeneous spatiotemporal response properties of GrCs^24^. Combining MFA inputs of opposite signs could therefore complement other circuit mechanisms^80,81,82^ that contribute to diversity in temporal properties of GrCs, which is essential for learning associations on different timescales.

Opposite sign response properties of different BPN-MFAs are also likely to contribute to the puzzling response properties of Golgi cells (GoCs), which receive sparse feedforward excitation from MFAs^83^. Local innervation of GoCs by PM or NM MFAs could drive the often anti-phase activity patterns exhibited by neighbouring cells^84^, which ride on top of a slower common mode modulation, that is thought to arise from the strong electrical coupling between these cells^85,86,87^. Innervation of GoCs by subgroups of MFAs with specific response properties could therefore generate diverse spatiotemporal patterns of inhibition onto GrCs, increasing their combinatorial diversity^80^.

While the BPN is the largest source of MFAs and projects widely across the cerebellar cortex^17,50,9^, some GrCs in Crus I/II will be partially or solely innervated by MFAs originating from other precerebellar nuclei. The extent of such mixing of input streams will depend on the density of innervation of the different MFA projections, which vary across regions^17,9,88^. Electrophysiological mapping of cutaneous receptive fields in CrusI/II (originating from the spinal trigeminal nucleus, Sp5^89,90^ revealed a ‘fractured map’ with similarly tuned responses in patches of a few hundred micrometres^91^. This suggests Sp5 inputs vary on spatial scales at, or larger than, our imaging volume. Our finding that BPN-MFB population activity lacks spatial structure on the 0-200 µm spatial scale, suggests spatially invariant mixing of cortico-pontine and other precerebellar MFA input to GrCs on local scales. Innervation of individual GrCs by MFAs from multiple precerebellar nuclei is likely to contribute to multimodality in GrCs^26^. This could also underlie the difference between the NM/PM ratio observed in GrCs^24^ (NM/PM = 0.3, and that predicted from our model with solely BPN input (0.47 - 0.59), if they had a greater preponderance of PM responses. Although multiple cellular mechanisms could contribute to the heterogeneity in GrC response properties, our results suggest that their bidirectional modulation is inherited, at least in part, from BPN-MFA inputs. BPN-MFA inputs are therefore likely to contribute to the nonlinear mixed population coding^25^ in GrCs that combines sensory, proprioceptive and motor information^24,36,38,92^.

### Implications for cerebellar granule cell layer function

Our results suggest that substantial numbers of BPN-MFAs are active during the QW and AS and that GrCs inherit their bidirectional population code^24^ from MFAs. Such a high level of excitatory drive to the cerebellar cortex is consistent with population imaging from GrCs, which have revealed that a large fraction of GrCs are active during behaviour^35,36^. Our results complement these findings by showing that BPN-MFA inputs are likely to be a major determinant of this dense GrC population activity. While this provides further evidence against the ultra-sparse coding envisioned by Marr and Albus^33,34^, theoretical studies suggest that denser high dimensional GrC population codes can maintain high information rates and perform pattern separation effectively^29,31,19^. Indeed, the optimal level of GrC population activity depends on multiple factors with denser codes providing better learning performance for time varying motor control^93^ and conservation of information^19^). The relatively high levels of activity we observe in the BPN-MFA population and those reported for GrC populations during spontaneous behaviours^24^, therefore suggest that the circuit is configured to deliver the high rates of sensorimotor information required for accurate motor control.

The GrC layer is thought to separate MFA activity patterns prior to associative learning in Purkinje cells. Theory and modelling predict that the sparse input connectivity of GrCs, their nonlinear thresholding and the much larger number of GrCs than MFAs perform nonlinear mixing, decorrelation, dimensionality expansion and noise reduction^33,34,30^. Consistent with this view, BPN-MFAs exhibit a high dimensional low redundancy population code during complex behaviours and near-random local mixing of MFAs with different response properties in the GrC layer. However, the high dimensional low redundancy BPN-MFA and GrC^24^ population codes we observe during complex behaviours differs from the redundant, lower dimensional code observed with simultaneous recordings of L5 pyramidal neurons in motor cortex and cerebellar GrCs during a skilled forelimb manipulandum task^38^. While it seems likely that the low dimensionality of the neocortical and cerebellar representations are limited by the low dimensionality of the forelimb task^94^, the high level of correlation between pyramidal neurons and GrCs and the lack of dimensionality expansion observed suggests the cerebellar input layer performs noise reduction rather than pattern separation under these conditions^38^. Although, further work is required to understand how GrC layer processing varies with behavioural complexity, our results show that signalling along the cortico-ponto-cerebellar pathway utilises a low redundancy, bidirectional, high dimensional population code during complex behaviours that is well suited for forming the nonlinear mixed, high dimensional GrC representation that have been measured^24^ and cerebellar theories have long predicted.

Another factor when considering cerebellar population codes is the ease with which sensorimotor information can be decoded and associations learned by downstream neurons. Electrophysiological experiments suggest that Purkinje cells can learn GrC activity patterns^95^ and operate in a largely linear regime^96^. Dense high dimensional GrC population codes can be learned more quickly than sparse codes using a Purkinje cell-like linear decoder^29,93^. Moreover, time-varying stimuli with temporal correlations encoded with bidirectional ON-OFF population codes can be linearly decoded more efficiently than ON-ON codes, reducing the number of spikes required for the same decoding performance^68^. However, learning in Purkinje cells is likely to be more complex than captured by simple linear decoders, since Purkinje cells in different microzones exhibit different mean firing rates and opposing response properties^97^. Interestingly, this added complexity can be easily accommodated, since our results suggest bidirectional MFA population activity propagates effectively through the cerebellar input layer. Selective innervation of ‘upbound’ and ‘downbound’ Purkinje Cells by PM or NM GrC axons, could therefore provide the excitatory drive for bidirectional learning across these populations^98^. While the details of implementation remain to be explored, our results show that the cortico-pontine input to the cerebellar cortex provides a rich sensorimotor basis set for downstream decoders to learn internal models of the animal’s state, an essential requirement for coordinating motor actions and predicting their sensory consequences^2,1^.

### Future directions

Our findings open many new avenues to explore. Key among these is whether the GrC layer increases the embedding dimensionality of sensorimotor representations and decorrelates inputs^30^ or whether noise reduction dominates^38^. Our finding that the response properties of BPN-MFAs are highly heterogeneous, both in space and time, raises the question of how temporal dispersion in responses is utilised by downstream neurons to form sensorimotor associations on different timescales. Key to understanding how the cerebellar cortex transforms and decodes the bidirectional population code will be to determine how PM and NM BPN-MFAs innervate Golgi cells and how opposite signed GrCs innervate upbound and downbound Purkinje cells. While addressing these points will provide a more comprehensive understanding of information processing in the cerebellar cortex, our current results provide key new insights into the population level properties of the major input to the cerebellar input layer.

## Acknowledgements

This project was supported by the Wellcome Trust (203048; 224499). R.A.S. is in receipt of a Wellcome Trust Principal Research Fellowship. SS was supported by the Wellcome Trust (225412/Z/22/Z). We acknowledge the GENIE Program and the Janelia Research Campus of Howard Hughes Medical Institute for making the GCaMP6 material available. We thank A. Hantman for providing the Slc17a7-Cre transgenic mice and N. A. Cayco-Gajic and H. Gurnani for advice on reusing their analysis scripts. We are grateful to Diccon Coyle and the SilverLab microscopy team for their technical support. We thank A. Barri, N.A Cayco-Gajic, D. Coyle, H. Gurnani, L. Justus, B. Marin, A. Valera and D. Whittaker for their comments on the manuscript.

## Author contributions

Conceptualisation: R.A.S. Methodology: H.R., Y.X., S.S. and R.A.S. Software: Y.X., S.S. Investigation: H.R. performed experiments. Analysis: H.R., Y.X., S.S. and R.A.S. Writing—original draft preparation: R.A.S. Writing—review and editing: H.R., Y.X., S.S. and

R.A.S. Supervision: R.A.S. Funding acquisition: R.A.S.

## Methods

### KEY RESOURCES TABLE

| REAGENT or RESOURCE | SOURCE | IDENTIFIER |
| --- | --- | --- |
| **Bacterial and virus strains** |  |  |
| AAV9-CAG-Flex-GCaMP6f-WPRE | Addgene | Cat # 100835-AAV9;<br>RRID:Addgene_100837 |
| **Chemicals, peptides, and recombinant proteins** |  |  |
| Red fluorescent beads | Thermofisher | Cat No. F8858 |
| **Deposited data** |  |  |
| Raw and analysed data | This paper | https://doi.org/10.6084/m9.figshare.28512758 |
| **Experimental models: Organisms/strains** |  |  |
| Slc17a7-IRES-Cre transgenic mice | Provided by Adam Hantman; generated by the Janelia Farm-Gene Targeting and Transgenics Facility | N/A |

| Software and algorithms |  |  |
| --- | --- | --- |
| Microscopy software | Nadella et al., 2016 | https://github.com/SilverLabUCL/SilverLab-Microscope-Software |
| Custom data-analysis scripts | This paper | https://github.com/SilverLabUCL/MossyFibres |
| Computational modeling and analysis | This paper | https://github.com/SilverLabUCL/MossyFibres |
| Custom scripts for preprocessing raw acquisition data | This paper | https://github.com/SilverLabUCL/MossyFibres |
| ImageJ (Fiji), Image processing software | Schindelin et al., 2012 | https://imagej.net/Fiji |

### RESOURCE AVAILABILITY

#### Lead contact

Further information and requests for resources and reagents should be directed to and will be fulfilled by the Lead Contact, Angus Silver.

#### Materials availability

This study did not generate new unique reagents.

#### Data availability

Data presented in the main figures and in Supplementary figures will be made available on FigShare: https://doi.org/10.6084/m9.figshare.28512758.

#### Code availability

The SilverLab LabVIEW Imaging Software is available on GitHub at: https://github.com/SilverLabUCL/SilverLab-Microscope-Software.

Analysis scripts are available at https://github.com/SilverLabUCL/MossyFibres

### EXPERIMENTAL MODEL AND SUBJECT DETAILS

#### Animal preparation for *in vivo* imaging

All experimental procedures were carried out in accordance with UCL Animal Welfare Ethical Review Body. All procedures were approved by the UK Home Office, subject to the restrictions and provisions contained in the Animals (Scientific Procedures) Act of 1986. Experiments were performed on 4 transgenic *Slc17a7-IRES-Cre* mice (on a C57B/6 background; 3 females, 1 male). All surgical procedures were carried out under sterile conditions, with separate procedures for viral delivery and cranial window preparation. After analgesic injection with buprenorphine (0.1 mg/kg), mice were deeply anaesthetised with a ketamine:xylazine mix (100:10 mg/kg) and mounted in a stereotaxic frame (Kopf Instruments). Then, 5 μl pipettes (BLAUBRAND, 7087-07) were pulled on a Sutter P97 micropipette puller, and suction filled with AAV9.CAG.Flex.GCaMP6f.WPRE.SV40 (Addgene Cat#100835-AAV9) to express genetically encoded calcium indicator GCaMP6f^99^ in the basal pontine nucleus (BPN).

Stereotaxic virus injections were performed on mice aged between postnatal day 50 and 80 (P50-P80). For viral delivery, stereotaxic coordinates were identified based on Allen Mouse Brain Atlas (BPN: (AP) 3.75 - 4.0 mm posterior to Bregma, (ML) 0.4 - 0.5 mm lateral to the midline, and (DV) 5.7 - 5.1 mm from the pia/dura). A small craniotomy was performed at 1-3 sites per animal (to maximise targeting efficiency), and a glass pipette (40-60 μm diameter) preloaded with the viral vector was slowly lowered into the identified coordinates. A pressure-based delivery system (Toohey Spritzer) was used to slowly inject 150-250 nL over 10-15 min into the BPN. In all animals this was accompanied by red fluorescent microbeads (4 μm fluospheres, ThermoFisher, 1:2000 dilution) that were injected in the cerebellum for real time 3D movement correction^51^. Analgesia (bupivacaine 0.05%) was then administered to the surgical wound site. After surgery, atipamezole (1 mg/kg) was administered for xylazine reversal. Post-operative care and analgesia (buprenorphine 0.1 mg/kg) and carprofen (5mg/kg) was administered for 48 hours.

### METHOD DETAILS

#### Headplate and cranial window surgery

After 2–6 weeks of AAV expression, a separate surgery was performed to prepare a cranial window for imaging. Prior to surgery, mice received subcutaneous injections of dexamethasone (1 mg/kg), atropine (0.04 mg/kg) and carprofen (5 mg/kg). Anaesthesia was induced by intraperitoneal injection of a mixture of fentanyl (0.075 mg/kg), medetomidine (0.75 mg/kg) and midazolam (7.5 mg/kg). Reflexes were monitored throughout the surgery, body temperature maintained at 37 °C using a regulated heating pad (FHC Inc.), and eyes covered by Viscotears eye gel to prevent dehydration. After removal of overlying skin, a custom head plate was centred above Crus I/II and attached to the skull using dental acrylic cement (Paladur, Kulzer). A 5 mm craniotomy was performed (with ∼1 mm durotomy) over the Crus I/II region, and the exposed cerebellar cortex was cleared with sterile cortex buffer (125 mM NaCl, 5 mM KCl, 10 mM glucose, 10 mM HEPES, 2 mM MgSO_4_, 2 mM CaCl_2_ (pH 7.4) to wash blood and remaining debris from the craniotomy. The craniotomy was then sealed with a 5 mm glass coverslip (630-2112, VWR) and fixed with cyanoacrylate glue to allow chronic imaging. Post-surgery analgesia (buprenorphine 0.1 mg/kg) was subcutaneously administered prior to anaesthesia reversal via atipamezole (3.75 mg/kg), flumazenil (0.75 mg/kg) and naloxone (1.8 mg/kg). Animals were kept in a heated chamber until full recovery of reflexes and locomotion, and provided post-operative care and analgesia (buprenorphine 0.1 mg/kg) and carprofen (5mg/kg) for 48 hours. Mice were group housed and kept on a 12-h light/dark cycle with food and water ad libitum. Mice were familiarised with the setup for 1–3 days and then imaged from 4 weeks after viral injection.

#### Behaviour and Data acquisition

##### Behaviour

Mice were free to stand or run on a cylindrical Styrofoam wheel. Forelimb and whisker movements together with running speed were monitored using a CCD camera at 300 Hz and 30 Hz respectively. Each imaging session consisted of 20 trials of 20 s each. For some sessions, a mild 50 ms air puff was delivered to the ipsilateral whiskers (non-aversive sensory stimulus) at 5 s within each trial. Behavioural recordings and air puff delivery was synchronised with imaging sessions by trigger signals.

#### 3D two-photon imaging of mossy fibre activity

Calcium imaging was performed in awake, head-fixed mice using a high speed custom-built Acousto Optic Lens (AOL) 3D two-photon microscope^49^ with real-time movement-corrected imaging^51^, that allowed measurement of many MFBs distributed within the 250 μm × 250 μm × 100 μm imaging volume. Following acquisition of a Z-stack of the imaging volume, regions of interest (ROIs) comprising multiple imaging patches and planes were selected at different depths to maximise the number of MFBs recorded. Imaging was performed at 920 nm with depth-dependent modulation of excitation intensity using a 20x water immersion objective (Olympus XLUMPlanFLN 20 × numerical aperture 1.0). Acquisition rates for different imaging sessions varied between 8-20 Hz depending on the number, size and type of ROIs. Before each recording session, the same imaging site was found by matching anatomical landmarks. Mice with bone regrowth under the window or poor viral expression were excluded from the study. For further details on the two-photon imaging configuration see ^24^.

##### Tissue preparation and confocal imaging

After in vivo imaging was complete, animals were deeply anaesthetised and their brains were fixed by transcardial perfusion with 4% paraformaldehyde in phosphate buffer 0.1 M followed by 24 h of postfixation in the same solution at 4 °C. The brains were then sliced sagittally. Slices (80 μm) were cut on a vibratome (VT1000S, Leica Microsystems) and washed in PBS. The slices were imaged with a confocal microscope (Leica TCS-SP8) and slide scanner (Zeiss AxioScan Z1).

##### Extraction of mossy fibre activity from imaging data

All analysis was performed using custom-written MATLAB scripts (version 2022a, Mathworks Inc.) or publicly available toolboxes. Imaging data for each patch were exported to TIFF files using in-house software written in LabVIEW (National Instruments). Before extracting fluorescence activity times series from patches, post hoc image registration was used to correct for any residual movement in the images^24,100^. Raw fluorescence time series F(t) were obtained for each MF bouton by averaging across pixels within each ROI using thresholding and particle analysis (FiJi). All pixels within the manually traced ROI mask were averaged to give raw MFB fluorescence. Baseline F and ΔF/F: The 10th percentile of raw fluorescence for each MFB was used as the baseline (𝐹*_b_*) and the normalised fluorescence activity (Δ𝐹/𝐹*_b_* ) was calculated as(𝐹 − 𝐹*_b_*)/𝐹*_b_*. For raw imaging data was stored as TIFF images for each time series. FiJi was used for 3D rendering and to adjust brightness and contrast of the images for display purposes. No deconvolution or nonlinear scaling such as gamma correction was used on the data for fluorescence traces.

##### Measurement of whisker position and locomotion

Two video cameras with far infrared LED illumination were used to monitor the face and whiskers. Facial areas were recorded at 1280 x 960 resolution at 30 Hz (The Imaging Source), whereas whisking was recorded at 644 x 484 resolution at 300 Hz (Mako). All behavioural data were acquired with the SilverLab custom software running under LabVIEW (National Instruments). Whisker pad movement was calculated using a previously-described measure of motion index^101^. The wheel motion index was calculated using a small ROI selected on the wheel, as the average difference in pixel values between successive frames^101^, smoothed over 200 ms. This provides an estimate of wheel motion without distinguishing between forward movement (for example, running) or backward movement (for example, startle responses). Datasets with no locomotion or whisking were excluded from the behavioural analysis as no comparison could be made with the representation of the active state in the same population of mossy fibres.

##### Behavioural states

A smoothed version of wheel MI and whisker MI (with a smoothing window of 100 ms) were used to calculate the behaviour states. Periods during which both the smoothed wheel MI and whisker MI simultaneously exceeded the threshold for a duration of more than 3 seconds were classified as the active state (AS). For periods less than 3 seconds were categorised as quiet wakefulness (QW). The threshold for each recording session was typically set as the 90^th^ percentile of wheel MI and whisker MI during the session. However, some manual adjustment was necessary for 3 sessions where the mouse was either highly active or very inactive.

#### Identification of boutons on the same mossy fibre axon

Mossy fibre boutons were grouped based on the similarity of their activity. Initial grouping was based on the activity of pairs of MFBs with the criteria being the 99^th^ percentile of pairwise Spearman correlation. Pairwise analysis was applied sequentially, comparing the first MFB activity to all others above it and then to the second MFB and above and so on. Each of these initial groups was then evaluated for their linear deviation ratio (LDR) with a criterion of <1.5^24^. After this automated procedure, a manual visual check was carried out to finalise the grouping of MFBs to correct any false positives and false negatives (7.7% and 1.6% respectively, *N* = 4). Groups were merged if they were visually identical or exhibited significant member overlap, defined as at least n−1 out of n members being common between the groups (n ≥ 3). In cases where the same MFB was assigned to more than one axon group, the average LDR of two groups was compared and the MFB was assigned to the group with the lower LDR. The activity of an MFA was calculated by averaging the ΔF/F across all its MFB members. To determine the distance between pairs of MFBs on an axon we selected the first MFB in the group as the reference point and calculated the Euclidean distance to all other MFBs in the group. The MFB with the shortest distance from the reference was identified as the nearest neighbour. This nearest point became the new reference MFB for the next iteration, until the inter-bouton distances from all MFBs within the group were calculated. This was repeated for all MFAs.

#### Identification of positively, negatively and not significantly modulated mossy fibre axons

MFAs that were positively modulated (PM), non-significant modulated (NSM), and negative modulated (NM) during AS when compared to QW were identified based on the significance of their correlation. To do this we calculated the Pearson correlation coefficient (𝐶𝐶_0_) between states, locomotion, and whisking and MFA activity data. Then, we randomly shuffled the behavioural data in time using blocks of 1s and obtained the shuffled correlation (𝐶𝐶_𝑠ℎ_) between the shuffled behaviour and neuronal activity. The process of obtaining the shuffled correlation was repeated 100 times. We computed the mean and standard deviation of these correlations and normalised the initial Pearson correlation coefficient using Z-score:

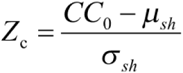

Here, 𝑍*_c_* is the normalised correlation coefficient. We defined 𝑍*_c_*> 2𝑆𝐷 as PM, 𝑍*_c_*<− 2𝑆𝐷 as NM, and 𝑍*_c_*< |2𝑆𝐷| as NSM.

#### Spatial distribution of PM, NM and NSM MFBs in the imaging volume

Spatial coordinates (X, Y, and Z) of MFBs across all mice (*N* = 4) were combined. The Kolmogorov-Smirnov test was used to evaluate whether the distributions of the X, Y, and Z coordinates for the PM and NM groups originated from the same distribution.

We evaluate the spatial distribution of MFBs using a 3D Ripley’s *K* and *L* function, which were developed to analyse spatial point patterns in three dimensions^59,105^. The Ripley’s *K* function, 𝐾(𝑟), quantifies the expected number of MFBs within a sphere of radius 𝑟 centred at a randomly chosen MFB, normalised by the overall point density:

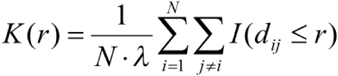

Where λ is the density of MFBs in the spatial domain, 𝐼(𝑑_𝑖𝑗_ ≤ 𝑟) is an indicator function that determines whether point 𝑗 is within a distance 𝑟 of point 𝑖.

Under the assumption of Complete Spatial Randomness (CSR), the points follow a homogeneous Poisson process. In this case, the expectation of Ripley’s *K* function in 3D space is given by:

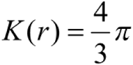

The Ripley’s *L* function is derived from Ripley’s *K* function:

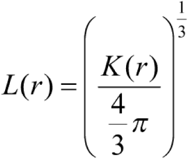

This transforms 𝐾(𝑟) into a function 𝐿(𝑟) that approximates 𝑟 under CSR.

#### Analysis of manifolds

The geometry of neuronal activity subspaces (manifolds) was analysed by performing PCA on the MFA population activity during QW and AS across concatenated sessions. The neural activity subspaces were visualised by plotting the top 3 principal components. The angle between the neural activity subspaces during AS and QW was assessed by measuring both the largest and smallest angles between the subspaces, using the first two principal components to define each subspace (for a detailed description see^24^). To determine the statistical significance of the observed angles between AS and QW state spaces, we randomly shuffled the time indices of the data in 10 s blocks and determined the principal angle between the first and second halves of these randomised data. In addition to comparing manifold angles, we also quantified the average Euclidean distance between data points. For each state, we calculated the intra-state (within manifolds) distances among all pairs of data points within that state. Similarly, inter-state (between manifolds) distances were measured between all pairs of data points across the two states.

#### Linear regression

To investigate how MFA activity was related to different behaviours, cross-validated linear regression (ridge) was used to predict locomotion, whisking, and states^24^. Two types of predictors were used: (1) principal components (PCs) extracted from MFA activity (Figure 5) and (2) activity traces of the MFAs (Figure 6).

##### PC-based decoding

For PC-based decoding, PCA was first performed on the neural activity of all MFAs from each session across the entire time period to reduce dimensionality and extract PCs (𝑋). A subset of the dataset from each recording session was selected for regression. Both the selected PCs (𝑋) and behavioural data (𝑌) were mean-centred by subtracting their respective means to ensure consistency.

For cross-validation, PCs data were separated into 80% for training and 20% for testing^24^.

Ridge regression was performed on the training dataset (𝑋_𝑡𝑟_) and training behaviour (𝑌_𝑡𝑟_) to obtain the regression coefficients (β), which were then applied to the test dataset (𝑋_𝑡𝑒_) to predict behaviour (𝑌):

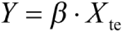

The explained variance (EV) quantifies the proportion of variance in measured behaviour (𝑌_𝑡𝑒_) explained by predicted behaviour (Ŷ) and was computed as:

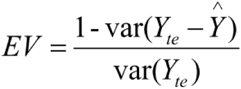

where 𝑣𝑎𝑟(𝑌 − Ŷ) represents the variance of the residuals. To obtain the cross-validated explained variance (CVEV), the EV values were averaged across all K folds (K = 5):

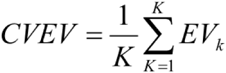

This process was repeated 10 times for each session to ensure robustness, and the final CVEV was obtained by averaging the results across repetitions.

##### MFA activity-based decoding

The similar ridge regression framework was applied in MFA activity-based decoding, with activity traces of the MFAs used as predictors (𝑋) and behaviour as response variable (𝑌). When examining the impact of MFA numbers on decoding performance, MFAs were not randomly selected as predictors. Instead, they were ranked using a combined score, which consisted of:

a. 50% from the Pearson correlation between MFA activity and behaviour, and
b. 50% from the ridge regression beta values obtained when using all MFAs as predictors.

Ridge regression was repeated by incrementally adding the top-ranked MFAs, and the corresponding CVEV was calculated for each subset size (Figure 6B). To evaluate the contributions of specific functional categories of MFAs (i.e. PM, NM, and NSM), we randomly selected MFA within each category, with the number of selected MFAs varying between 1 and 30. Ridge regression was then performed for each subset, and the decoding performance was assessed in the same way as described previously (Figure 6D).

##### Determine optimal number of PCs or MFAs

To determine the optimal number of PCs, we used an early stopping criterion: if the improvement in CVEV was less than 0.01 for five consecutive increments, the number of PCs corresponding to the start of this sequence was selected as the optimal value (Figure 5D).

To determine the minimum number of MFAs required to predict behaviour, we applied lasso regression with penalties ranging from λ = 0.001 to 0.05. The minimum number of non-zero coefficients was recorded for each penalty. The decoding performance of the optimal number of MFA was compared to that of the single best MFA using the lasso regression (Figure 6C).

#### Dimensionality Analysis

We used a bicross-validated version of PCA to infer the dimensionality of neural representations during spontaneous behaviour^57,104^. To control for differing population sizes across experiments we randomly subsampled 100 MFAs for each session (Method details^24^). For each session, we randomly selected 80% of the data and designated it as training data (𝑡𝑟), while 20% of the data was designated as testing data (𝑡𝑒). We perform singular value decomposition (SVD) to decompose (zero-centered) neural population activity into orthogonal population modes. We calculated the first k PCs from the training dataset (with a default block size of 1 second), resulting in the following low-rank approximation:

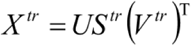

where 𝑈 are the loadings, 𝑆 is a diagonal matrix of covariance along different modes (eigenvalues of covariance matrix), 𝑉 is the dynamics along different modes and 𝑇 is the transpose.

To cross-validate these PCs we further split the test data into a second partition of training neurons (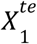 80 % of the population) and test neurons (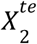, 20% of the population). The low-rank matrix decomposition for this test data subset can be expressed in block format as:

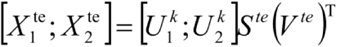

where 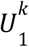 and 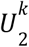 represent the PC directions of 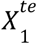 and 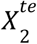, respectively. We used the upper block to estimate the 𝑘 latent dynamics (*S*^𝑡𝑒^*V*^𝑡𝑒^) via linear regression and use the lower block to predict 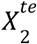. After learning the low-dimensional subspace from the training data, the testing data was projected into the same subspace. We predicted the activity of the test neurons from 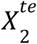 in the low-dimensional subspace:

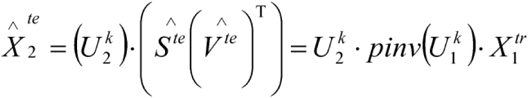

The subsampling was repeated 10 times for each recording session. In each iteration, the minimum number of principal components needed to maximize the explained variance of 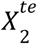 was determined. The lower bound of the dimensionality is the number of PCs required to maximize the explained variance of 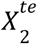. The extrapolated dimensionality, representing the number of PCs required to achieve 100% variance explained, was estimated using linear extrapolation across experiments.

To quantify and interpret the impact of subsample size on the variance and corresponding dimensionality, we computed a slope based on the maximum variance (𝑋) and corresponding dimensionality (𝐷):

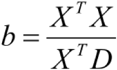

MFA per dimension was calculated as:

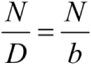

#### Modelling

We based our GrC layer connectivity model on the finding that each GrC receives synaptic input from 4 different MFAs^106^, one onto each of its short dendrites^22^ (21.7 μm). We used the experimentally measured positions of the PM, NM, or NSM MFBs from an experiment with a large population of MFAs. We then placed 300 GrCs randomly within a subvolume 21.7 μm from the maximum and minimum X, Y, and Z positions of the outer boundaries of the volume occupied by the MFBs. We calculated the pairwise distances from each GrC to MFBs and selected the 4 MFBs with the closest distances as the ones most likely to be connected to the GC. To ensure GrCs were innovated by 4 different MFAs, any MFBs originating from the same axon were excluded and replaced with the next closest MFB based on pairwise distance rankings. We recorded the input types (PM, NM and NSM) of the four selected MFBs for each GrC and calculated the average probability of occurrence for the 15 different MFA input combinations across 10 repetitions of the model.

To evaluate how our experimentally constrained GrC model compared to purely random sampling we calculated the theoretical distribution for 4 random samples of 3 object classes without replacement. This corresponds to a hypergeometric distribution. The probability of each combination of PM, NM, and NSM connections was calculated as follows:

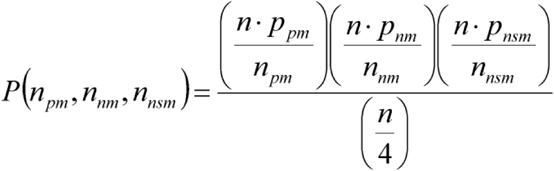

where, 𝑃_𝑝𝑚_,
𝑃_𝑛𝑚_,
𝑃_𝑛𝑠𝑚_ are the fractions of PM, NM, and NSM inputs, respectively and 𝑛_𝑝𝑚_,
𝑛_𝑛𝑚_,
𝑛_𝑛𝑠𝑚_ are the number of each input type within a given combination of four MFBs.

To predict the response properties of GrCs in the GrC layer connectivity model, each type of MFB was assigned a synaptic strength (1 or 0) during a specific state. PM = 1 during the AS and 0 during QW state. NM = 1 during the QW state and 0 during AS. NSM = 0 for both AS and QW states. The GrC response was determined by summing the synaptic input weights from the four MFBs as the state changed from QW to AS. For example, an increase in synaptic weight from QW to AS corresponds to a positively modulated GrC. To evaluate the effect of different GrC spike thresholds, different response thresholds were applied, where GrCs were only considered responsive if their activity changed between QW and AS in a positive or negative direction and their inputs summed to a value above threshold (Th = 0,1,2,3) in one or both of the states. GrCs with subthreshold or constant summed synaptic input weight across states were considered NSM.

#### Statistical tests

All statistical tests were two tailed. Data are presented as mean ± s.e.m unless stated otherwise. Throughout the manuscript, *n* refers to the number of experiments, and *N* refers to the number of animals. All error bars on figures indicate s.e.m. For paired data, we used the Wilcoxon signed-rank test. To compare distributions, we used the Kolmogorov-Smirnov test, while group means were compared with the Mann-Whitney U test and considered significant when p < 0.05. All correlations are reported as the Pearson correlation coefficient (𝐶𝐶_𝑝_) and Spearman correlation coefficient (𝐶𝐶_𝑠_) used in axon grouping.

---

**Supplementary Figure 1.**
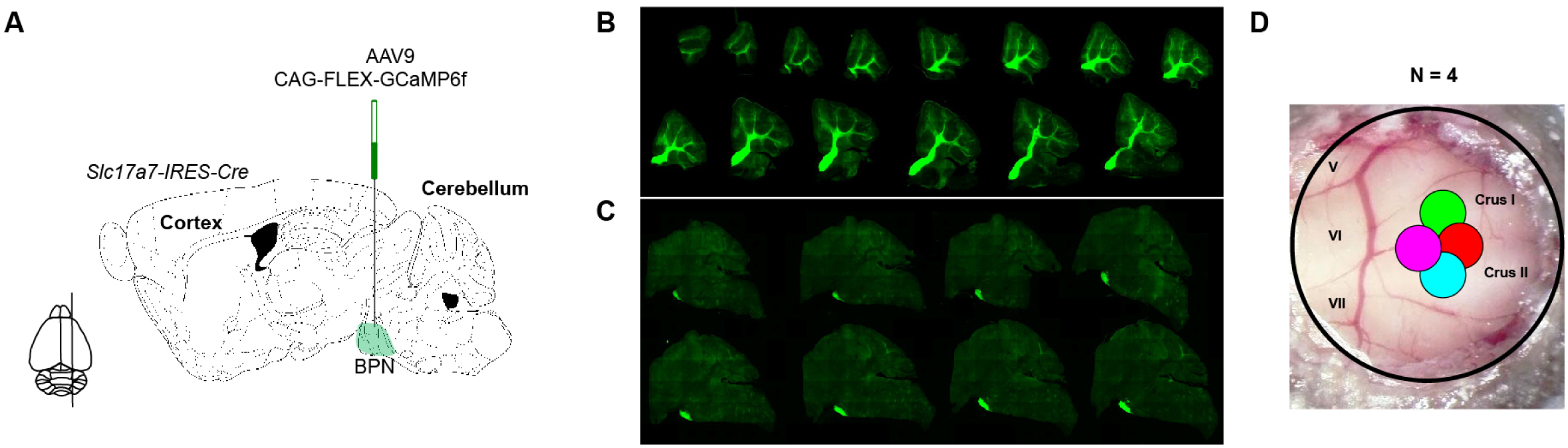
**Anatomical confirmation of injection and recording sites** (A) Schematic diagram showing viral injections of AAV9-Flex-GCaMP6f construct into basal pontine nucleus (BPN) that used the CAG promoter to drive targeted expression in Slc17a7-IRES-Cre transgenic mice. (Modified from Paxinos and Watson Mouse Atlas). (B) GCaMP6f fluorescence in fixed sagittal sections of cerebellar cortex, showing widespread expression in the granule cell layer of cerebellar Crus I/II. (C) GCaMP6f fluorescence localised to BPN injection sites. (D) Location of imaging of BPN mossy fibre axons in Crus I/II of the cerebellar cortex (N = 4 mice; magenta, cyan, red, green).

**Supplementary Figure 2.**
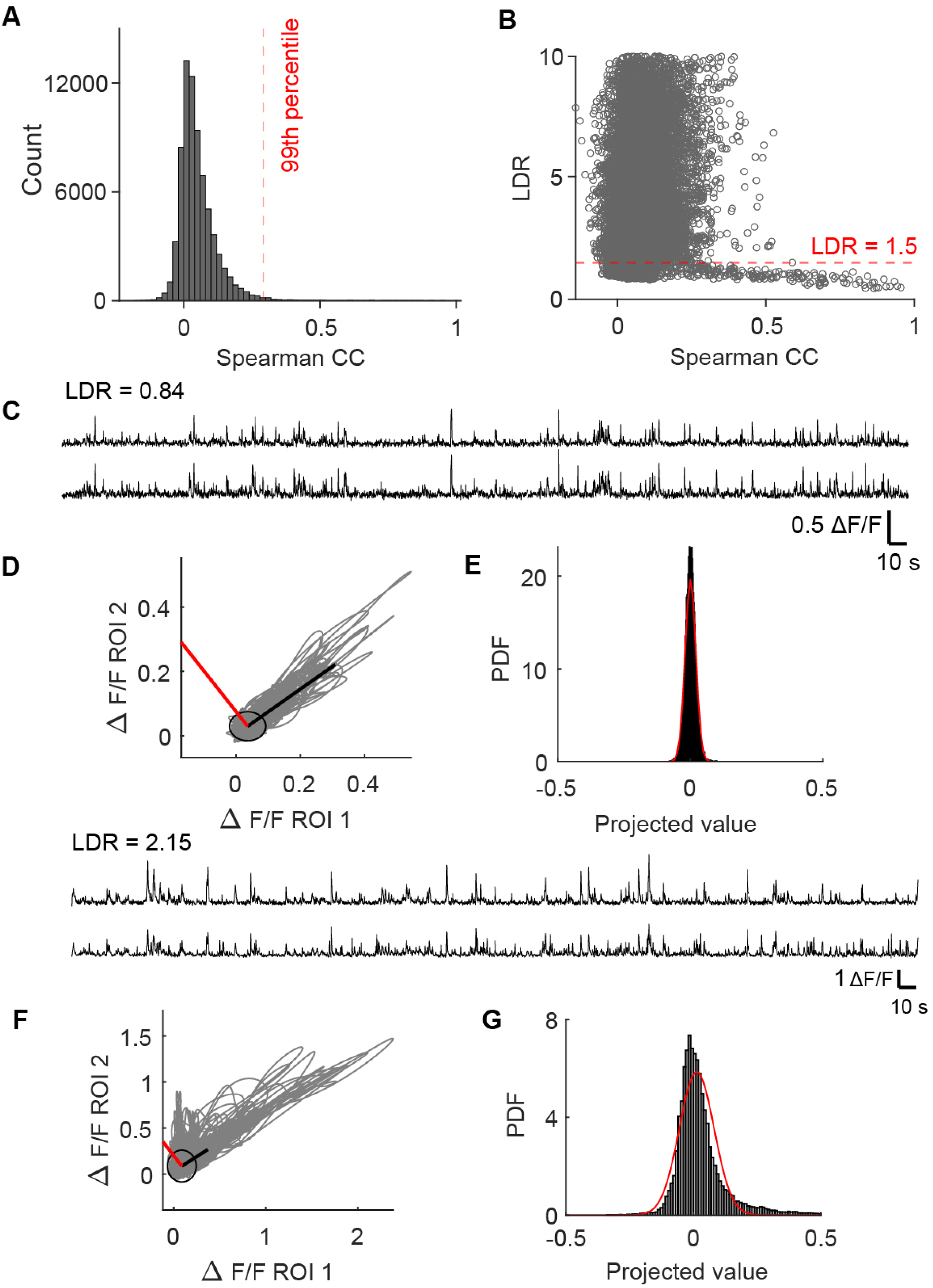
**Identification of boutons associated with the same axon using activity correlation and linear deviation ratio criteria** (A) Distribution of Spearman pairwise correlations (CCs) of activity from one recording session. Dashed red line shows the 99th percentile. (B) The relationship between the Linear Deviation Ratio (LDR) and CCs of activity in (A). Red dashed line shows the 1.5 LDR threshold criterion. (C) An example of two mossy fibre bouton (MFB) activity traces with relatively high Spearman correlation (CCs = 0.48) that also passed the LDR threshold. (D) Activity of trace 1 from (C) plotted against another activity trace. The black line indicates a fit from linear regression. The red line indicates the vector onto which activity is projected to calculate the LDR. (E) Histogram of activity from (D) projected onto the vector (black), and the analytically calculated distribution of the baseline distribution projected onto the vector (red curve). The ratio of the variances of these distributions is used to compare with the LDR threshold (LDR = 0.83). (F)-(H) Same as (C)-(E) for a pair of MFB activity traces with a relatively high Spearman correlation (CCs = 0.51) that failed to pass the LDR threshold (LDR = 1.9). Red arrows in (F) indicate parts of the traces that are clearly different between the two traces.

**Supplementary Figure 3.**
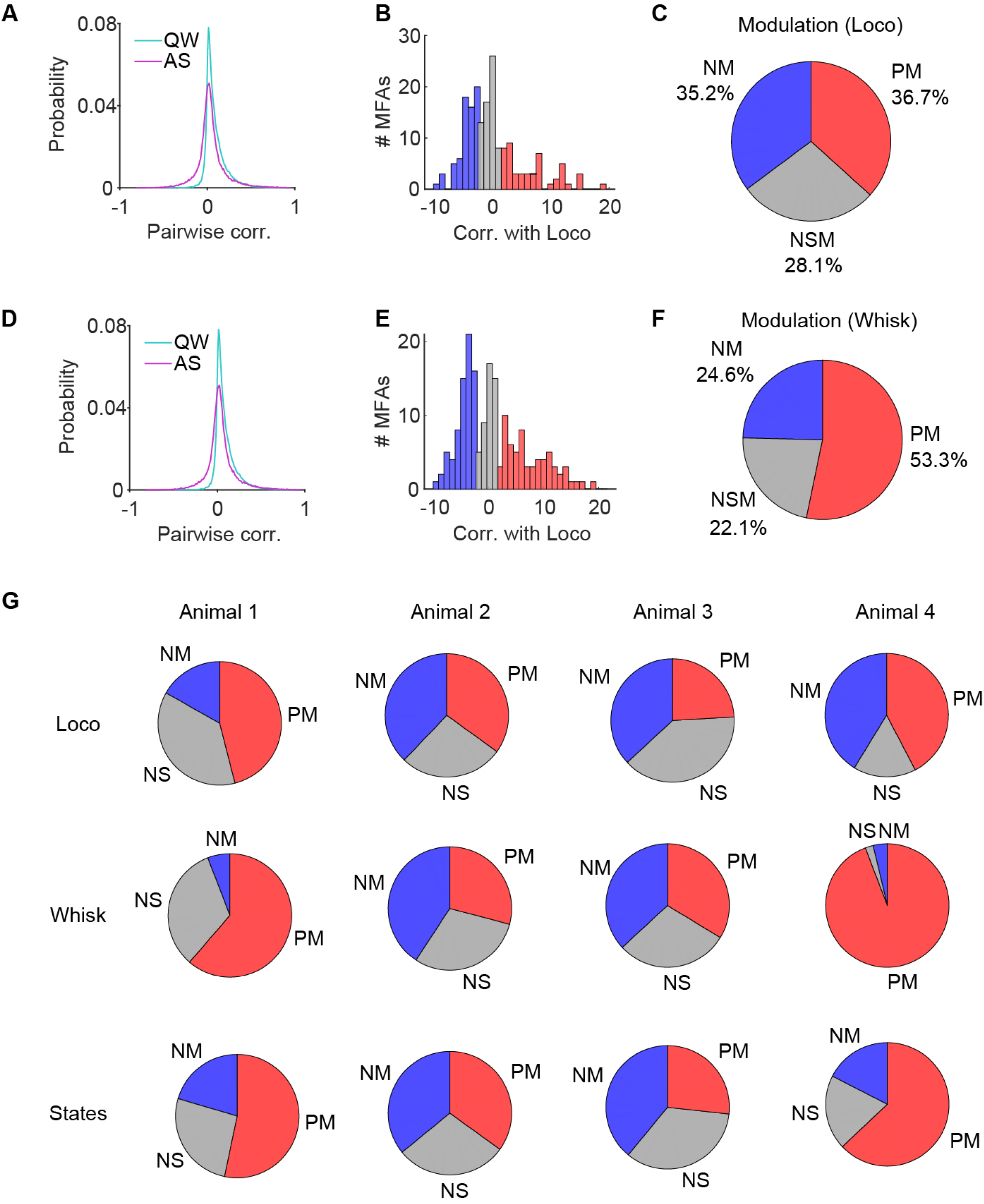
**PM/NM/NSM fractions for individual animals** (A) Correlation of the z-scored activity for MFAs and locomotion. The black histogram shows the correlations that are significant (> 2SD) (B) Normalised correlations for z-scored activity for MFAs exhibiting significant positive modulation (PM, red), negative modulation (NM, blue) and non-significant modulation (NSM, grey) for locomotion. (C) Fractions of significant PM, NM and NSM across animals (n = 16, N = 4). The level of significance was P <0.05. (D)-(F), same as (A)-(C), but shows the correlation of the z-scored activity for MFAs and whisking. (G) , Fractions of significant PM, NM and NSM for each animal across different behaviours (n = 16, N = 4).

**Supplementary Fig. 4.**
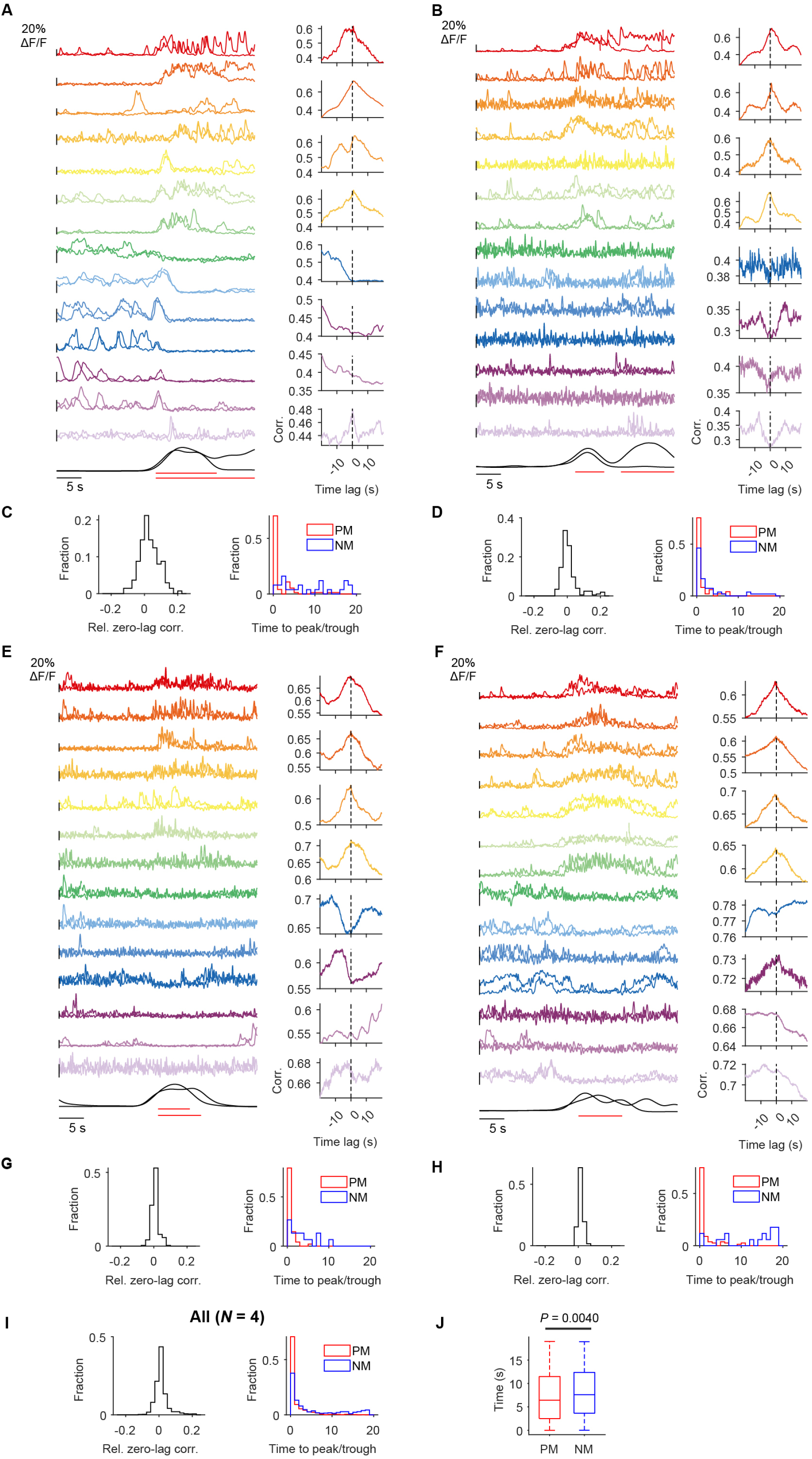
**Reliability of mossy fibre axon activity** (A)(B)(E)(F) Sample activity (ΔF/F) of 14 MFAs 20s before the onset of running and after the onset of running for two separate episodes of running for four animals (left to right). Running speed is shown on the bottom for each episode. Activity traces are colour coded according to positive modulation (top) or negative modulation (bottom) of activity with locomotion. Plots to the right of the traces show normalised cross-correlograms of MFA activity with running for sample MFAs. Colour code same as in (A). (C)(D)(G)(H) Left: Distribution of relative zero-lag correlations for all MFAs for each animal. The relative zero-lag correlation is calculated from the cross-correlograms in (A), as the difference of the cross-correlation at lag zero (l=0) and the average cross-correlation preceding it (from l = −10 to 0). Right: Distribution of the time to peak/trough of the cross-correlation for positively-modulated (PM) and negatively-modulated (NM) MFAs across animals. Time to peak is calculated as the lag at which the maximum cross-correlation appears after zero lag (from l=0 to l=20s) for PM MFAs. Time to trough was calculated as the lag at which the minimum cross-correlation appears after zero lag (from l=0 to l=20s) for NM MFAs. (I) as above but for all animals. (J) Peak times for PM and trough times for NM are calculated based on significant increases or decreases (Δ > 3 STD) across all sessions, with time = 0s marking the onset of running. Mann-Whitney U test.

**Supplementary Figure 5.**
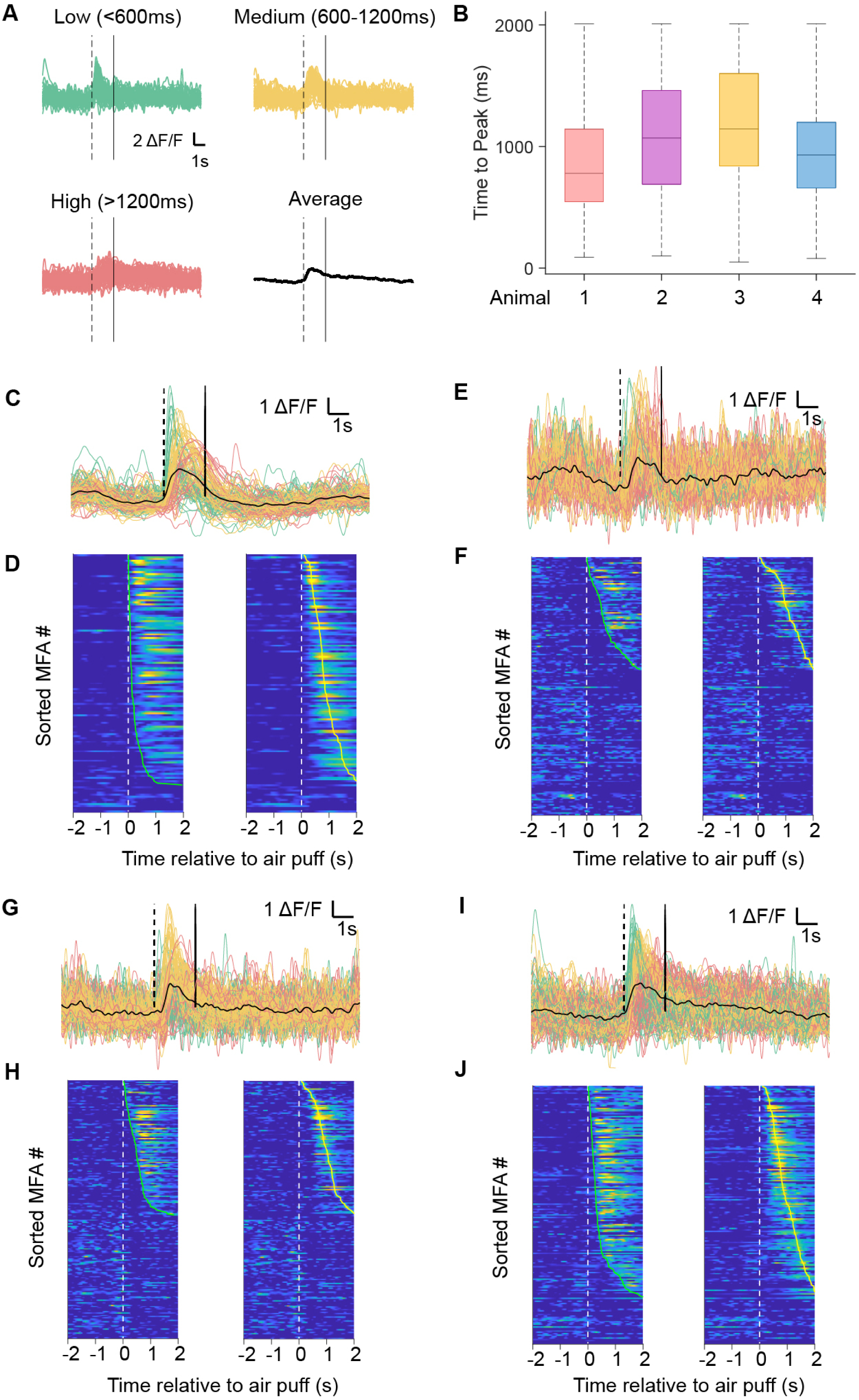
**Reliable and heterogenous air puff-evoked axon responses** (A) Mossy fibre axon (MFA) responses to air puff for Animal 1 across trials. Latency was categorised into three groups based on time delays following air puff delivery: <600 ms (low latency, green), 600–1200 ms (medium latency, yellow), and >1200 ms (high latency, red). Latency was calculated as the time difference between air puff onset (dashed line) and the time point where a significant increase in ΔF/F activity is observed (>3 SD above baseline). The onset of activity is defined as the first point showing a significant increase that leads to a peak. (B) Summary of time to peak for each animal. The data includes all sessions (n = 16 sessions), with mean and standard deviation shown in the below table. (C)(E)(G)(I) ΔF/F activity traces for all trials of an example session from each animal. Each trace represents one trial (20 trials per session). Vertical dashed lines indicate air puff onset at 5 s, and vertical solid lines mark the 20s observation window used to assess significant increases in ΔF/F activity. (D)(F)(H)(J) Heatmaps of averaged MFA ΔF/F activity for an example session from each animal (Animals 1 - 4). Each heat map is sorted according to increasing latency of responses relative to puff onset (t = 0; left: onset time, right: peak time). These results confirm air puff-evoked MFA responses are reliable from trial-to-trial and that there is a large degree of heterogeneity in the latency of MFA responses across the local population.

**Supplementary Figure 6.**
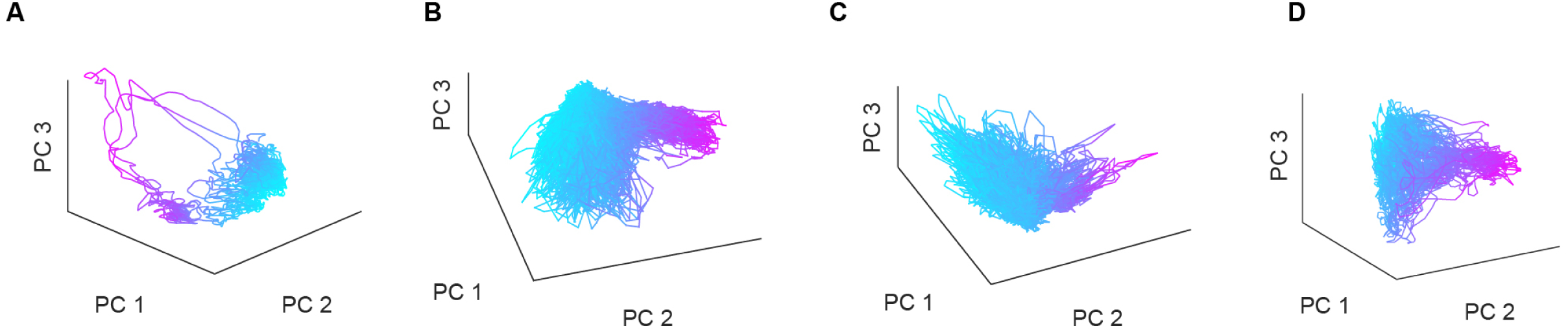
**Activity state manifolds across behavioural states for individual animals** (A) -(D) The first three principal components (PC) of mossy fibre population activity for each mouse. Colours as in Figure 3.

**Supplementary Figure 7.**
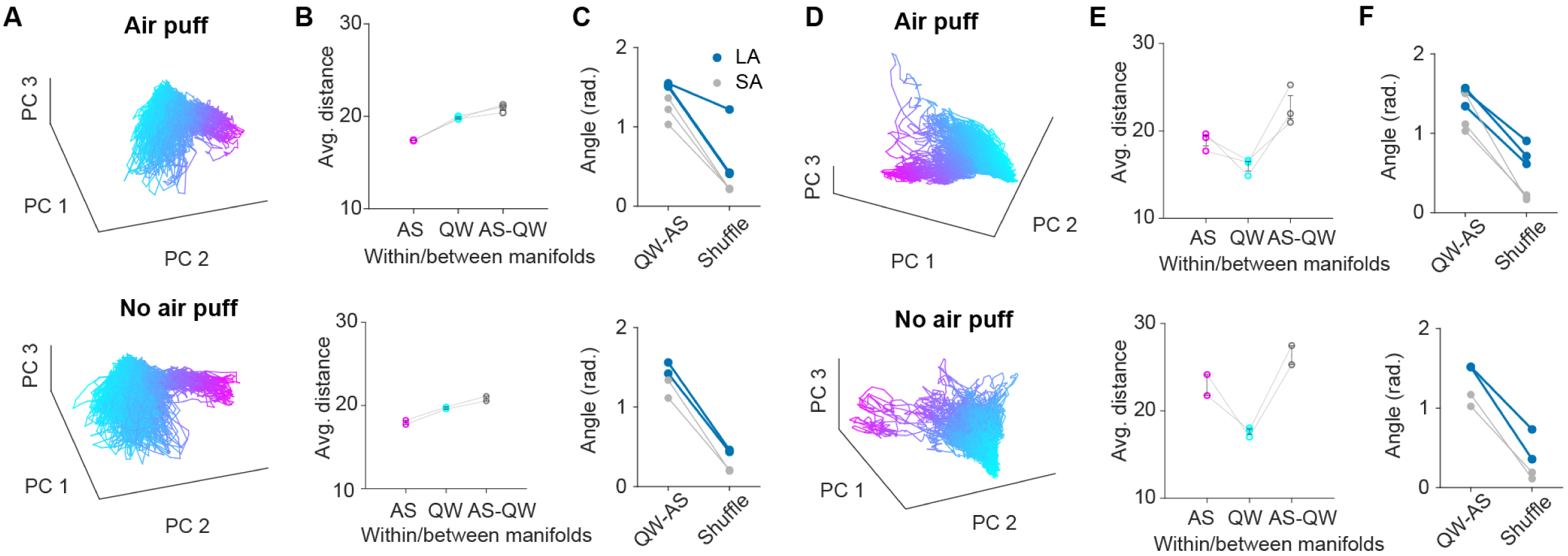
**Manifold analysis across behavioural states, with and without air puff** (A)(D) The first three principal component (PC) trajectories of BPN-MFA population activity for Animal 2 (A) and Animal 4 (D), with (n = 2 for Animal 2, n = 3 for Animal 4) and without an air puff (n = 3 for Animal 2, n = 2 for Animal 4). (B)(E) The average Euclidean distance between all pairs of neural activity patterns during different behavioural states (ΔF/F) within the AS manifold (AS-AS), within the QW manifold (QW-QW), and between the two manifolds (AS-QW). (C)(F) Smallest and largest angles between the QW and AS manifolds in the same experiment, along with the null distribution, obtained by calculating the angle between two halves of the data, after temporally shuffling (t = 10s) each experiment for Animal 2 (C) and Animal 4 (F), with or without an air puff.

